# Exposure to lysed bacteria can promote or inhibit growth of neighbouring live bacteria depending on local abiotic conditions

**DOI:** 10.1101/2020.10.23.352005

**Authors:** Fokko Smakman, Alex R. Hall

## Abstract

Microbial death is extremely common in nature, yet the ecological role of dead bacteria is unclear. Dead cells are assumed to provide nutrients to surrounding microbes, but may also affect them in other ways. We found adding lysate prepared from dead bacteria to cultures of *E. coli* in nutrient-rich conditions suppressed their final population density. This is in stark contrast with the notion that the primary role of dead cells is nutritional, although we also observed this type of effect when we added dead bacteria to cultures that were not supplied with other nutrients. We only observed the growth-suppressive effect of dead bacteria after they had undergone significant lysis, suggesting a key role for cellular contents released during lysis. Transcriptomic analysis revealed changes in gene expression in response to dead cells in growing populations, particularly in genes involved in motility. This was supported by experiments with genetic knockouts and copy-number manipulation. Because lysis is commonplace in natural and clinical settings, the growth-suppressive effect of dead cells we describe here may be a widespread and previously unrecognized constraint on bacterial population growth.

## Introduction

In any natural environment, microorganisms face environmental stressors that can kill them. These include abiotic factors like temperature, nutrient or pH stress, and biotic factors such as infectious viruses (bacteriophages) and antibiotics produced by other microorganisms. Because cell death is expected to release compounds such as amino acids (1), it potentially enhances growth of surrounding cells (2,3). Indeed, exposure to dead cells has been shown to support growth of neighbouring live bacteria (4,5) and is thought to play a key role in population survival of bacteria during extended periods after other nutrients are exhausted (1,3,6–8). However, dead cells and associated debris may also induce other types of changes in gene expression in living cells through the release of, for example, eDNA (9), membrane vesicles (10,11) or signalling molecules (12,13). For instance, in *Pseudomonas aeruginosa*, cells lysed by a competing strain cause neighbouring live cells to upregulate a number of genes involved in antibacterial activity, improving their fitness in interspecific competition (14). It was recently shown that dead cells can alter swarming behavior in *E. coli* and induce changes in gene expression that benefit swarming *E. coli* when challenged with antibiotics (15). These findings suggests the ecological role of dead cells may not be limited to nutrition (recycling nutrients and other compounds beneficial for growth).

Much research on the ecological role of dead cells has focused on experimental populations of bacteria that have entered late stationary phase or death phase in batch culture, and subsequent recycling of dead cells present in such cultures as nutrients (1,3,7,16–18). For example, *Escherichia coli* can grow in the supernatant of starved cultures, and the amount of growth depends on the density of the starved culture (3). Although such studies reveal the effects of changes in medium composition caused by bacterial growth, starvation and death, they do not tell us about the effects of dead cells themselves, because they are typically filtered out of the medium before the supernatant is added to live cultures. A recent study overcame this limitation by adding UV-killed culture directly to live cultures, demonstrating a nutritional effect consistent with that suggested by supernatant experiments (8). However, key gaps in our knowledge remain. Perhaps most importantly, it is unclear whether exposure to dead cells has a similarly nutritious effect in populations that are also supplied with other nutrients (because nutrient cycling effects are typically measured in starved cultures or nutrient-free medium). Bacteria in nature are not always starving, and responses to dead cells may be different when cells have access to other nutrients. Additionally, the impact of dead cells may depend on how they have died. It has been shown that live cells respond differently to dead kin depending on the cause of death in unicellular algae (19), yeast (20), and the eukaryotic immune system (21,22). Bacteria killed by stressors other than starvation may therefore have different effects on living cells. In particular, cell lysis, which is common in nature as a result of bacteriophage infection (23), some antibiotics (24,25) or Type VI secretion systems (26), is likely to release a greater fraction of intracellular contents into the environment and may therefore have different effects compared to those observed for starved cultures.

Here, we investigate the effects of various types of dead bacteria and associated cell debris/lysate, on population growth of *E. coli*. We do this in both nutrient-rich and nutrient-poor conditions, revealing environment-dependent effects: we show dead bacteria can serve as nutrients when these are scarce but can, surprisingly, negatively affect final population density in seasonal environments where other nutrients are supplied and then exhausted. By resuspending cultures in fresh medium immediately before killing them, we disentangle the effects of dead bacteria (cell debris and released compounds) from medium conditioning during prior growth. We do this for cells killed in various ways, such as heat, sonication and phage lysis. We then use transcriptomic analysis and genetic knockouts to show that dead bacteria trigger widespread changes in gene expression in neighbouring live cells, indicating a key role of upregulated motility genes in the observed population-level response to dead cells in nutrient-rich conditions. Together, these results suggest dead bacteria can have qualitatively different effects on neighbouring live cells depending on the local abiotic conditions and how they have died.

## Results

### Exposure to cell lysate can inhibit or increase population growth depending on abiotic conditions

To test the effect of dead bacteria, we resuspended cultures of *E. coli* in fresh medium immediately before killing them by sonication, after which we filtered out any surviving cells (our control treatment here was sterile medium subjected to the exact same procedure; see methods). Adding this dead-cell preparation (lysate) to live *E. coli* in buffer solution with no other carbon source (M9) supported increased population growth relative to the control treatment: the change in bacterial abundance over 24h estimated by optical density was significantly higher in cultures supplemented with dead cells (Welch two-sample *t*-test: *t* = −46.33, df = 4.51, *p* < 0.0001; Fig. 1). By contrast, when we added dead cells (lysate) to *E. coli* cultures in nutrient-rich medium (LB), we observed a reduction of final population density compared to cultures without added dead cells (Welch two-sample *t*-test: t = 15.79, df = 6.68, p < 0.0001, Fig. 1). When we estimated bacterial densities by plating and counting colony-forming units (CFUs) instead of optical density (OD), we observed a similar pattern (Fig. 1), with a ~7-fold increase in mean density in M9 (mean ±.s.d. = 2.28×10^7^ ±2.79×10^7^ CFU/mL without, 1.60×10^8^ ±3.47×10^7^ CFU/mL with dead cells) and a ~2-fold decrease in LB (mean ±s.d. = 1.58×10^9^ ±2.71×10^8^ without, 7.18 ×10^8^ ±4.02×10^8^ with dead cells). In a separate control experiment, we accounted for any possible nutrient depletion in the dead-cell preparation after resuspension and prior to sonication and filtration. We did this by including an additional control treatment where we resuspended *E. coli* as in the dead-cell treatment, but did not sonicate prior to filtration. This showed no effect on both OD and CFU (methods, Fig. S1), whereas both of these measures of population growth were negatively affected by addition of sonicated-and-filtered dead-cell preparation in the same experiment (Fig. S1). This indicates the negative response we observed in our dead-cell treatments in nutrient-rich medium was not due to nutrient depletion during dead-cell preparation. We also found the magnitude of the response to dead cells was stronger when the initial amount of sonicated cells was higher (148.5μl instead of 50μl of killed-cell suspension, with the same total culture volume of 150μl and live-cell inoculum of 1.5μl overnight culture; see Methods) in both rich medium (dead cells × amount interaction in LB: *F*_1,20_ = 18.93, *p* < 0.001; Fig S2) and in buffer (dead cells × amount interaction in M9: *F*_1,20_ = 41.87, *p* < 0.0001; Fig S2), although the negative effect in LB was observed for both ratios (Fig. S2).

**Figure 1.**
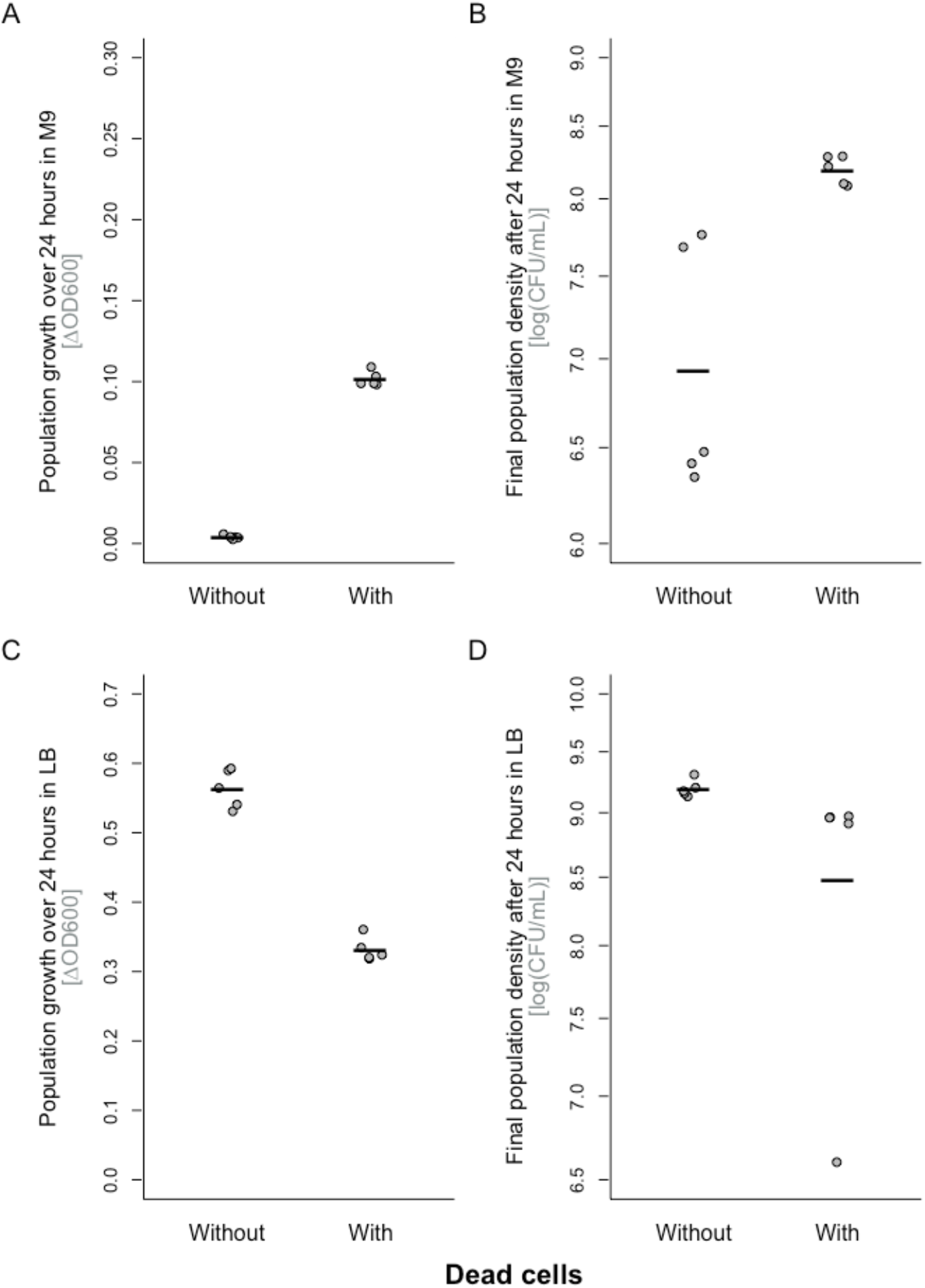
Population growth of *E. coli* K12 MG1655 in minimal medium (M9 without carbon source; A & B) and nutrient-rich medium (LB; C & D), in the presence and absence of dead-cell lysate (*x*-axis; lysate produced by sonication and filtration). Population growth is shown by two different measures: (A&C) change in optical density (OD600) over 24 hours and (B&D) colony-forming units per milliliter (CFU/mL) of the same cultures. All points represent independent replicates (*n* = 5) and individually prepared cultures of dead cells/controls. Each line shows the mean for one treatment group. We used a total volume of 150 μL, consisting of 1.5 μL overnight culture with 148.5 μL of dead cell suspension.

When we added cells lysed by bacteriophage T7, instead of by sonication, to cultures of a closely related, T7-resistant *E. coli* strain in nutrient-rich medium, we observed a similar negative effect as above for sonicated cells (Welch two-sample *t*-test, *t* = −4.72, df = 3.09, *p* = 0.017; Fig. S3). This indicates the negative effect of dead cells (lysate) we observed above for sonicated cells also applies to cells lysed by natural causes, including bacteriophages.

By contrast, we found cells killed by heat did not support significant growth when added in M9 (Welch two-sample *t*-test, *t* = 2.17, df = 9.96, *p* = 0.055; Fig. S4A), and had a very minor suppressive effect on OD in LB (Welch two-sample *t*-test for the effect of adding 50 μL dead cells, *t* = 2.39, df = 13.35, *p* = 0.032; Fig. S4B). Sonicating the cell suspension before heat-treatment restored their negative effect on population growth (Welch two-sample *t*-test: for OD *t* = −12.45, df = 7.25, *p* < 0.0001; for CFU/mL *t* = −12.45, df = 7.25, *p* < 0.0001, Fig. S5) in LB. Microscopic observations of different dead-cell suspensions showed those that were only heat-treated contained large clumps of intact, non-viable cells, whereas the phage-lysed suspensions and those that were sonicated before heat-treatment did not (Fig. S6). Thus, the dead-cell preparations that had the strongest negative effects on population growth of neighbouring live cells above were those associated with relatively extensive cell lysis and release of intracellular material.

### Adding more nutrients results in stronger growth suppression in response to dead cells

We hypothesized that the different responses to dead-cell preparations we observed in nutrient-rich medium compared to buffer solution are linked to the different amounts of population growth supported in the two environments. As a first manipulation of this, we tested if the response to dead-cell treatment (lysate) changed when we added lower amount of nutrients in LB (by using lower concentrations of tryptone and yeast extract, but keeping the salt concentration constant). The negative effect of our dead-cell treatment declined with decreasing nutrient concentration (dead cells × nutrient concentration interaction: *F*_1,44_ = 155.96, *p* < 0.0001; Fig. 2), eventually switching to a positive effect similar to that observed in M9 at the lowest nutrient concentration. When we added dead-cell preparation to cultures growing in buffer supplemented with other types of carbon sources, we observed variable, predominantly positive effects on growth (dead cells × medium interaction: *F*_3,48_ = 6.96, *p* < 0.001; Fig S7), similar to those observed in reduced-nutrient-LB treatments supporting equivalent final population densities. Thus, the growth-suppressive effect of exposure to dead-cell lysate on the final population density of live cultures was strongest in nutrient-rich conditions supporting a lot of population growth.

**Figure 2.**
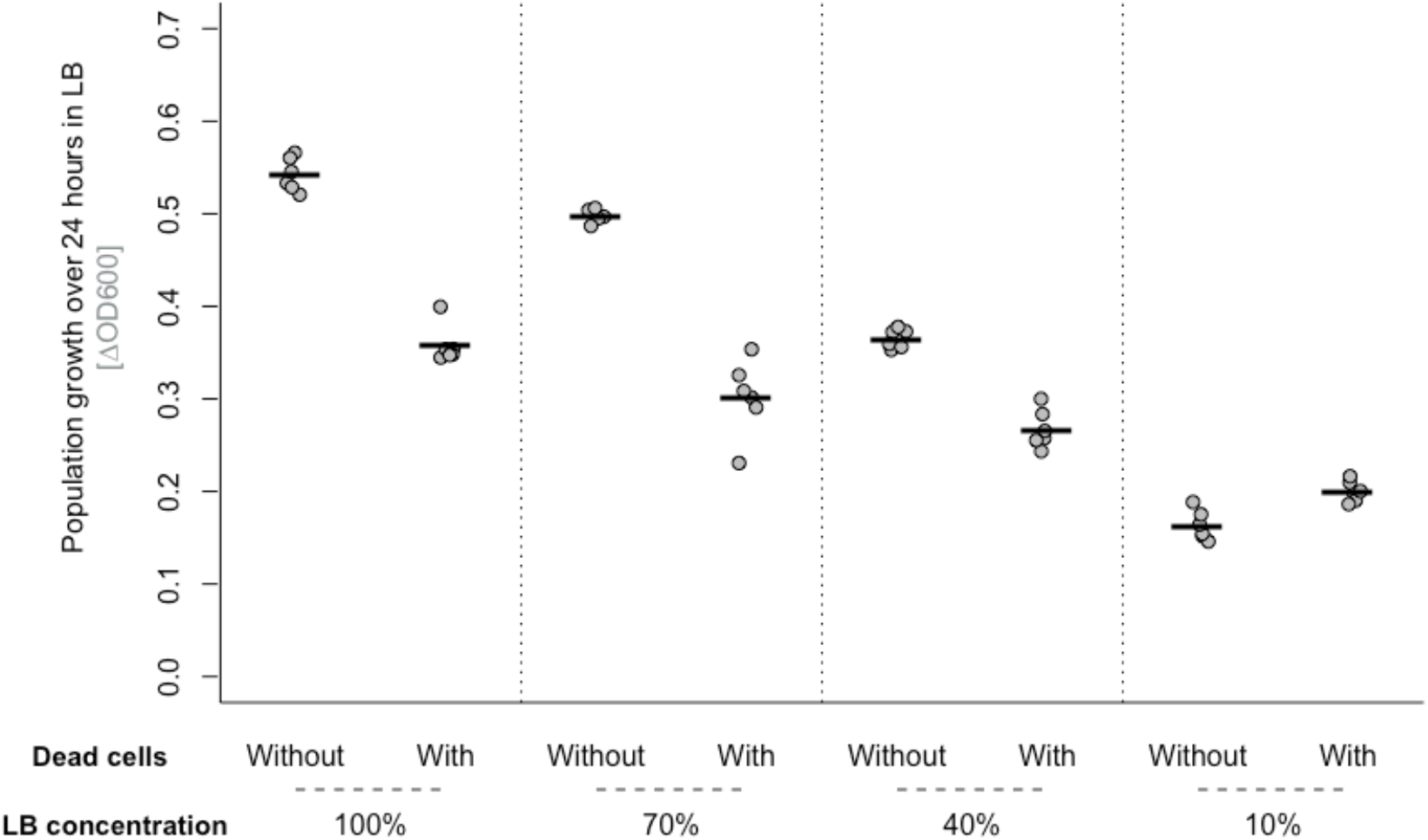
Bacterial population growth (change in OD600 over 24 hours) in the presence and absence of dead-cell lysate or control medium, in culture medium with different concentrations of the principle nutrients in LB (tryptone and yeast extract). Lower-nutrient-concentration treatments were prepared with 5 g/L NaCl, to retain the same osmolarity as in 100% LB. All points represent independent replicates (n = 6). Each line shows the mean from one treatment group. We used a total volume of 150 μL, consisting of 1.5 μL overnight culture with 148.5 μL of dead cell suspension (produced by sonication and filtration).

### Dead cells have similar effects in cultures with different population densities

The variable effects of dead-cell preparations above could potentially be explained by density dependence (exposure to dead bacteria having stronger negative effects per capita when the live population is at higher density, as is reached in higher-nutrient LB treatments). This could come about by, for example, interference with density-dependent behaviors such as those mediated by quorum sensing (27,28). Alternatively, dead bacteria could inhibit population growth in nutrient-rich medium by reducing the number of new cells produced per unit resource, with a per capita effect independent of the density of the culture. This would be more consistent with mechanisms such as nutrient sequestration or changes in gene expression that affect the efficiency of growth and replication. To disentangle these possibilities, we varied the starting density of the live *E. coli* population. Within the range tested, this did not significantly alter the net change in population density in the absence of dead-cell treatment in rich medium (effect of starting density in LB: for OD *F*_2,12_ = 2.98, *p* = 0.089, Fig 3; for CFU *F*_2,12_ = 2.18, *p* = 0.16, Fig. S8). In the presence of dead-cell treatment (lysate produced by sonication and filtration), we observed the same reduced population growth in LB as in our previous experiments above, and the magnitude of this effect was independent of starting population density (dead cells × starting density interaction: for OD, *F*_2,24_ = 1.67, *p* = 0.21, Fig. 3; for CFU, *F*_2,24_ = 0.93, *p* = 0.41, Fig S8). Thus, the growth suppression we observed was not stronger for cultures growing at higher densities, and therefore was probably not caused by interference with density-dependent growth regulation mechanisms such as quorum sensing. Consistent with the effect of dead bacteria depending more on growth dynamics than on population density, we also found the positive effect of dead bacteria on population growth in M9 was independent of the starting population density of the live bacteria (dead cells × starting density interaction: for OD *F*_2,24_ = 1.18, *p* = 0.32; for CFU *F*_2,24_ = 0.04, *p* = 0.96; Fig S9).

**Figure 3.**
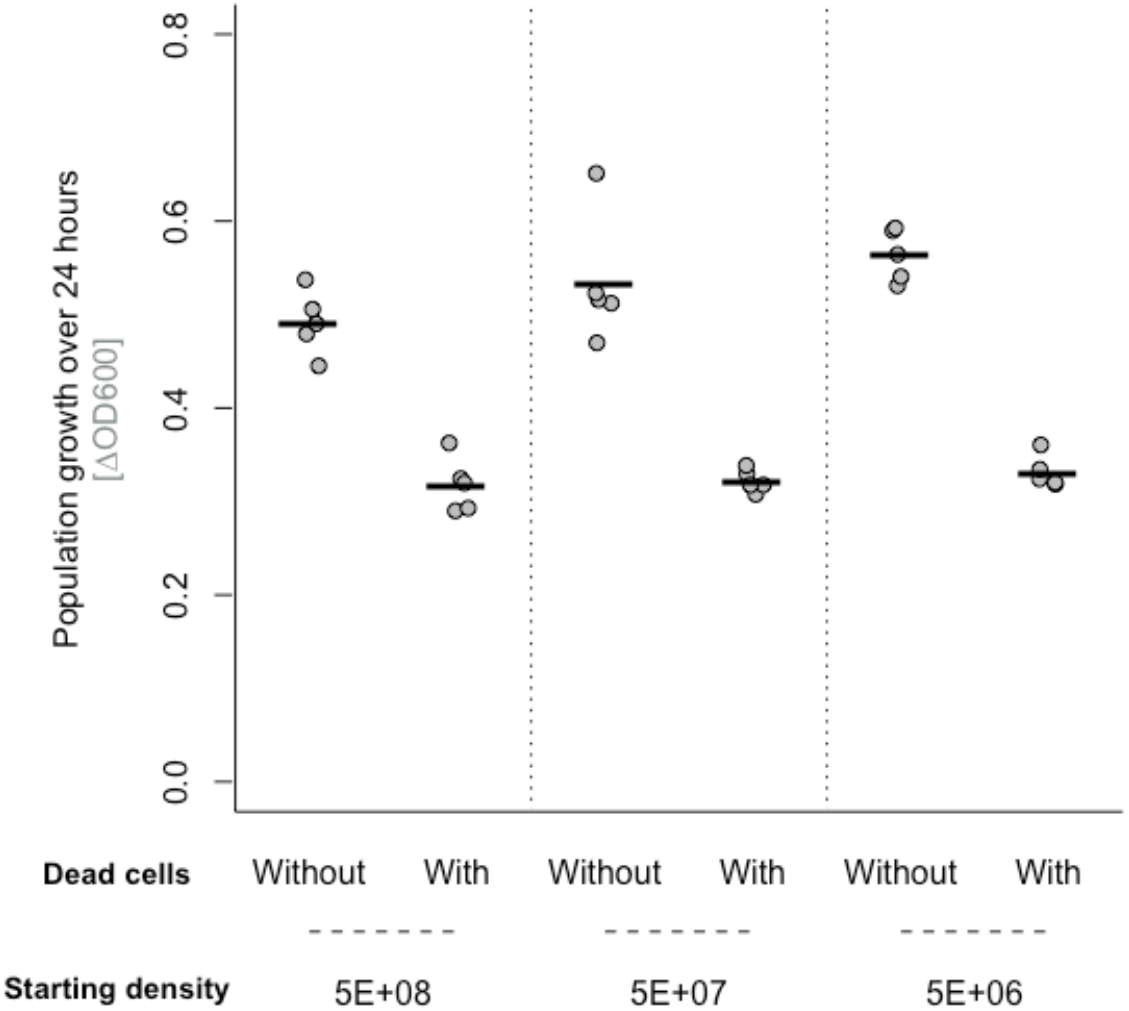
Bacterial population growth (change in OD600 over 24 hours) in nutrient-rich medium (LB) in the presence or absence of dead-cell lysate or control medium, using different starting densities of the live cells. In all other assays, 5e+06 was used as the starting density. All points represent independent replicates (n = 5). Black lines show the mean. We used a total volume of 150 μL, consisting of 1.5 μL overnight culture with 148.5 μL of dead cell suspension (produced by sonication and filtration).

### Exposure to dead bacteria induces changes in gene transcription in *E. coli*

We next aimed to find out whether the negative effect of dead bacteria on population growth of *E. coli* in rich medium (LB) was associated with changes in gene expression in the live cell population. We performed RNA sequencing on *E. coli* populations grown with or without dead cells (sonicated-and-filtered dead-cell preparation) to identify genes involved. We found a total of 321 genes to be differentially expressed in the dead-cell treatment compared to the dead-cell-free treatment (average expression change of at least two-fold and false discovery rate<0.1; see methods; Fig. 4, Table S1). 53 of these genes were differentially expressed at more than one of the three timepoints we sampled, of which 32 were exclusively upregulated. Of these upregulated genes, 19 were involved in motility, chemotaxis and flagellum synthesis (Fig. 4; Table S1). This was further corroborated by enrichment analysis using gene ontology (GO) categories, which showed significant enrichment of a total of 11 GO categories in the upregulated genes (table S2). Of these eleven categories, four were motility-associated, including two that were enriched at more than one timepoint. Individual motility genes that were upregulated included multiple genes of the *flg* (9) and *fli* (7) operons, as well as *motA, fliA* and *flhC*. These three genes are all transcriptional activators of motility and chemotaxis in *E. coli* (29–31), and therefore probably contribute to the observed increased expression of other motility/chemotaxis-associated genes, which is consistent with recent work demonstrating that the presence of dead cells can affect *E. coli* swarms and increase their antibiotic resistance. We also observed a shift in expression for several genes involved in purine (*pur* operon) and pyrimidine (c*arA* and *carB*) biosynthesis, ribose uptake systems (*rbsABCD)*, and the ferric citrate (Fec) uptake system, including *fecIR, exbBD, tonB* and the *fep* operon.

**Figure 4.**
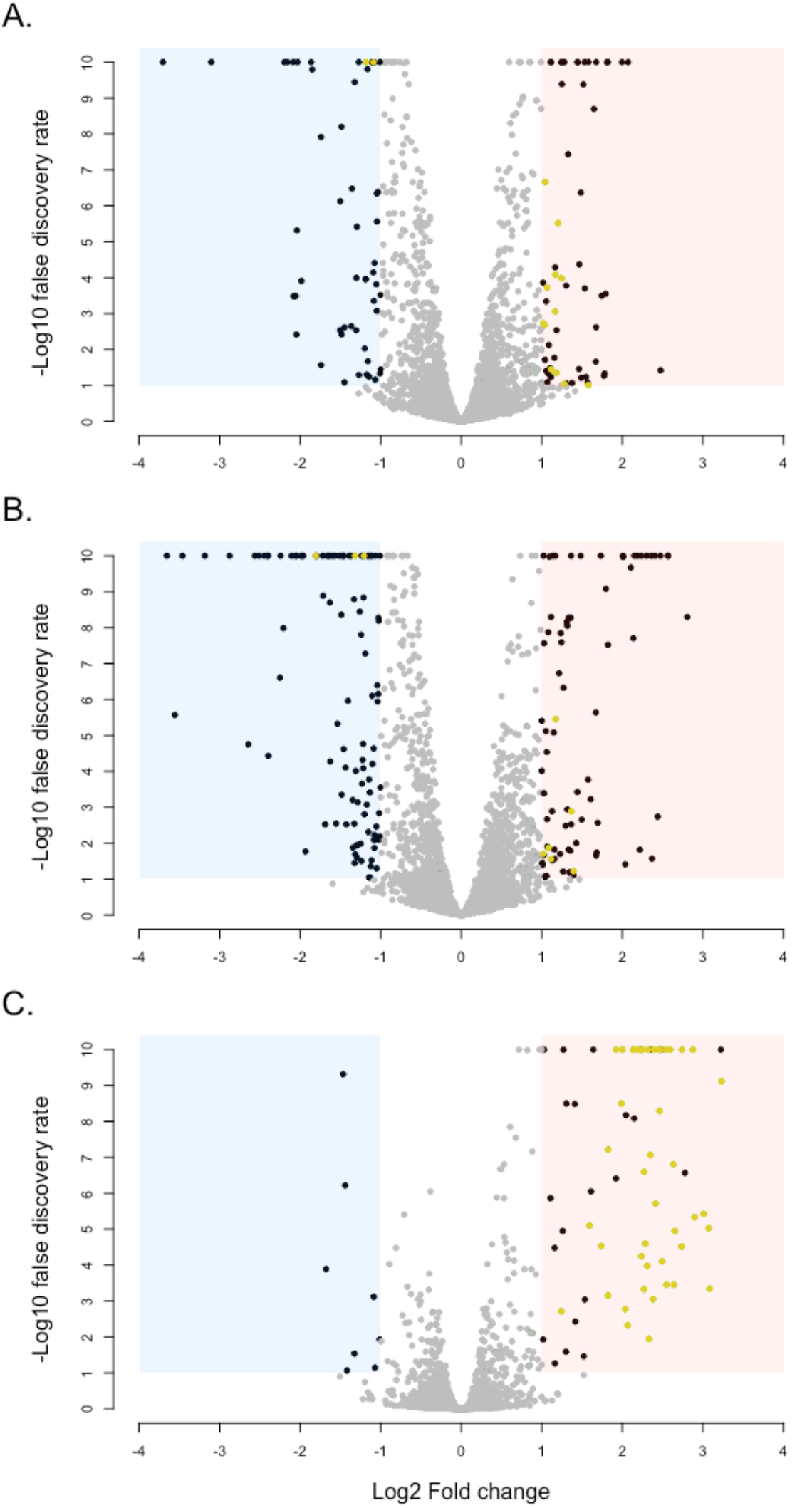
Transcriptional changes of *E. coli* in the presence vs absence of dead-cell lysate. Average gene expression relative to the control treatment is shown after (A) 5, (B) 6.5 and (C) 24 hours of growth in the presence of dead cell lysate (produced by sonication and filtration) in LB medium. Each point shows one gene. Dark points in the red and blue areas show significantly differentially expressed genes (false discovery rate<0.1 and fold change>2); yellow points show differentially expressed genes that are motility-associated (defined here as falling under one of the following five GOterms: GO:0071973 (bacterial-type flagellum-dependent cell motility); GO:0071978 (bacterial-type flagellum-dependent swarming motility); GO:0044780 (bacterial-type flagellum assembly); GO:0006935 (chemotaxis); GO:0044781 (bacterial-type flagellum organization)).

### Altered expression of motility genes affects population growth and response to dead cells

To test for further evidence that changes in gene expression identified above were involved in responding to dead cells, we assayed single-gene knockout strains in the presence and absence of dead cells. We did this for nine genes, focusing on those with regulatory functions and showing consistent fold changes across time points above, either themselves or downstream (Materials & Methods), using the same protocol and ratio of dead:live cells as above in our RNA sequencing experiment. We found the response to dead cells varied among different knockout strains (strain × dead cells interaction, *F*_8,72_ = 2.704, *p* = 0.01; Fig 5). Most knockout strains showed a reduction in final population density in the presence of dead cells, as observed for the wild type here and in our other experiments. However, both the *fliA* and *flhC* knockout strain did not respond to dead cells (Welch two-sample *t*-test - *fliA*: *t* = 0.61, df = 7.05, *p* = 0.56; *flhC*: *t =* −0.96, df = 6.25, *p* = 0.37; Fig. 5), consistent with them being involved in the wild-type response to dead cells. A later repetition of this assay with the higher dead:live cell ratio used in some of our other experiments confirmed these mutants responded more weakly than the wild-type to dead cells (strain × dead cells interaction, *F*_3,86_ = 4.24, *p* = 0.007; Fig. S10). *flhC* is one half of the *flhCD* dual transcription regulator which, among other genes, controls expression of *fliA. fliA* encodes σ^28^, a minor alternative sigma factor involved in regulation of genes involved in motility, chemotaxis and flagellar synthesis (29–31). Many genes downstream of *flhC* and *fliA* were also upregulated in our transcriptomic analysis (Fig. 4; Table S1). Despite this, we found no consistent effect of exposure to lysed cells on swimming motility of the wild type on soft agar (Fig. S11). We also found no evidence of altered biofilm formation, which can be affected by motility genes (34,35), in response to dead cells in LB (Fig. S12). The lack of detectable changes in swimming behaviour could be because the observed overproduction of transcripts involved in chemotaxis does not translate to an altered motility phenotype, but we do not exclude that were altered motility phenotypes not detected by this type of assay, particularly given there are several different types of motility in *E. coli* (36,37), and recent evidence that exposure to dead cells induces altered swarming behaviour upon antibiotic exposure (15).

**Figure 5.**
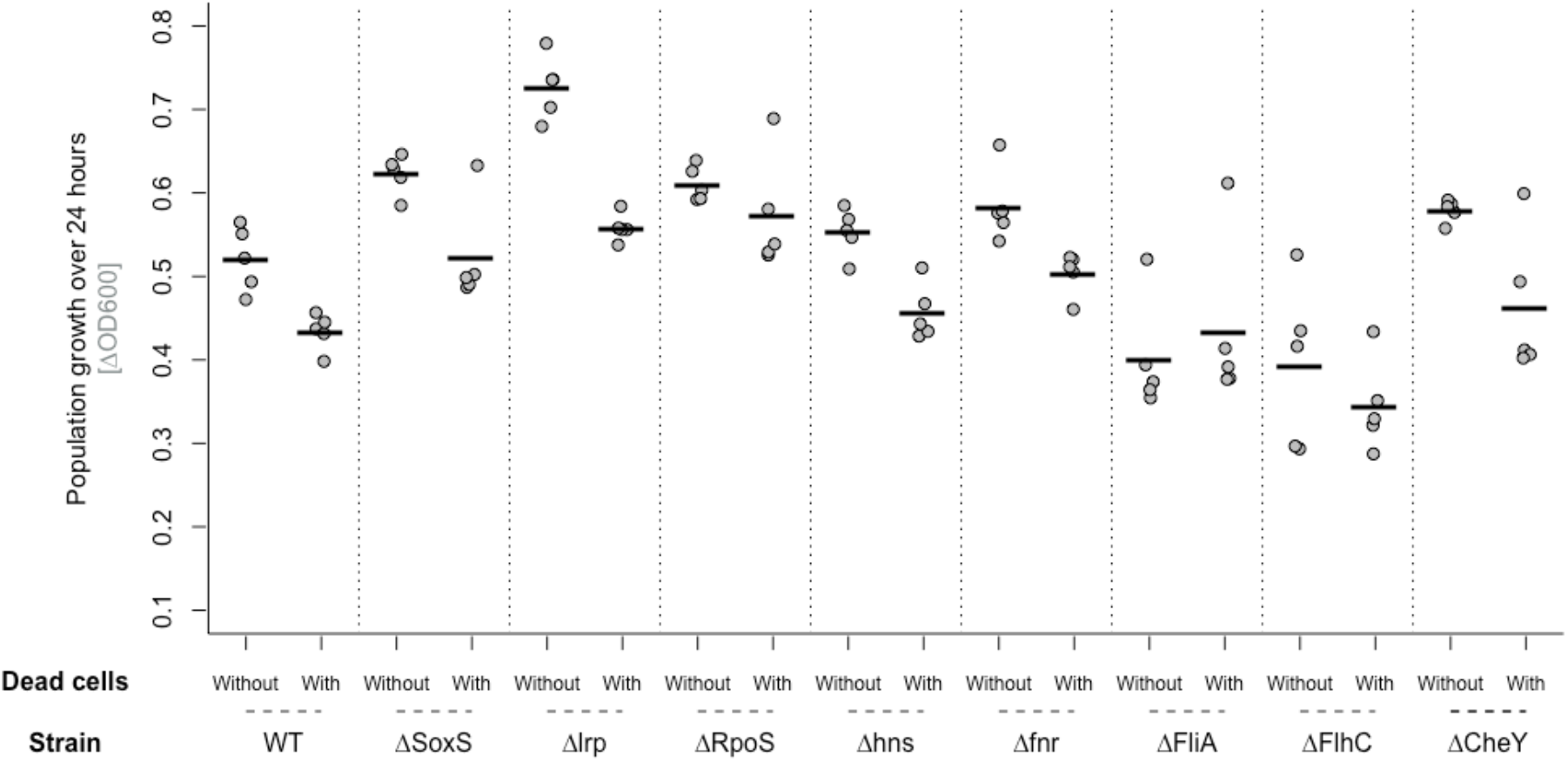
Bacterial growth (change in OD600 over 24 hours) of nine single-gene deletion mutants of *E. coli* in the presence and absence of dead-cell lysate (50μL sonicated-and-filtered lysate added to 100μl culture) in LB medium. All points represent independent replicates (*n* = 5). The line shows the mean. We used a total volume of 150 μL, consisting of 1.5 μL overnight culture with 148.5 μL of dead cell suspension.

Having observed that the two motility genes *fliA* and *flhC* (and genes they regulate) were overexpressed in the presence of dead cells, and that deleting these two genes reduced the negative response of *E. coli* to dead cells, we hypothesized that overexpression of these genes would negatively affect bacterial growth, which would help to explain the reduced growth we observed in the presence of dead cells. We tested this using ASKA library (overexpression) strains for *flhC* and *fliA*, which contain a high copy-number plasmid with the gene of interest under control of the *lacZ* promoter, inducible by IPTG. We found adding IPTG caused a significant reduction in bacterial growth for both the *flhC* and *fliA* overexpression strains (Fig. S13), consistent with overexpression of these genes being costly. By contrast, adding IPTG did not change average growth of K12 MG1655 in the same experiment (Fig. S13), suggesting the observed growth reduction for the ASKA strains did not reflect a general effect of IPTG on bacterial growth. When we added dead cells, neither *flhC* nor *fliA* overexpression strains showed reduced growth (Fig. S13). In the presence of IPTG, this may be explained by them already paying the cost of overexpression of these genes in both the presence and absence of dead cells. In the absence of IPTG this is more surprising, because we would expect them to show reduced growth in response to dead cells as in our other experiments. This may be explained by strain differences between the ASKA host K-12 AG 1 and K12 MG1655 (used in our other experiments, and showing the same reduced growth in response to dead cells as before when tested in the same block of assays as the ASKA strains; Fig. S12), or by leaky expression of the plasmid-borne gene copies in the ASKA strains (38). Consistent with the latter, the ASKA strains reached lower stationary phase densities than K12 MG1655 even without induction by IPTG.

## Discussion

We investigated the effect of dead bacteria on live bacteria. We found that in low-nutrient environments, adding lysate prepared from dead bacteria increased the final population density of neighbouring live cells, consistent with the widely held view that dead cells play a nutritional role. By contrast, we found that in a seasonal environment where bacteria were supplied with other nutrients, adding dead-cell lysate negatively affected final population density. The type of dead cells was important: we found a large effect using phage-lysed or sonicated cells, whereas cells that were killed by heat showed a marginal effect (note the lack of effect for heat-killed cells was not due to heat-lability of active ingredients; sonication prior to heating restored the negative effect on live cells). Our transcriptomic study of the response of *E. coli* to dead bacteria in LB revealed upregulation of flagellar synthesis and motility-associated genes. Knocking these genes out supported their involvement in the response to dead bacteria. By contrast, overexpression of the same genes was costly to bacteria in terms of population growth in these conditions, which probably contributes to the reduced population density we observed upon exposure to dead-cell lysate. Extensive cell lysis is common in nature, for example by phages (23), antibiotics (24,25) or type VI secretion (26), indicating such effects may apply in conditions beyond those tested here.

Together, these data show while dead cells can serve as nutrients, they can also promote other behavior in bacteria that reduces net population growth. This is in contrast to much previous work, which has often focused on the role of dead cells in nutrient recycling in starving cultures (3,5,8,39,40), late stationary phase and so-called GASP (Growth Advantage in Stationary Phase) phenotypes (1,6,7,18,41). In line with this research, we observe that *E. coli* can indeed grow on dead cells in buffer with no other available carbon source. However, when supplied with other nutrients, we see that this positive effect becomes progressively smaller the more nutrients are present, and eventually becomes negative. These different observations potentially reflect the different physiological state of *E. coli* in starvation/with no other nutrients supplied compared to in nutrient-rich medium. Induction of the stringent response via RpoS (42), and the subsequent wide-ranging changes in gene expression, may be an important determinant of the ability of *E. coli* to feed off dead cells in these conditions. This is corroborated by the central role of RpoS mutations in the GASP phenotype that appear in late stationary phases, and presumably confer increased ability to use nutrients released by dead cells (7). Another difference between our experiments and work with starved cultures is how cells have died (lysis vs starvation). Cause of death has been shown to influence responses to dead cells in several other systems (19–22). Thus, we do not rule out that starved cells, which are commonly used to study nutrient recycling of dead cell material, may have different effects to those observed here for lysed cells because, for example, prolonged starvation may cause cells to break down intracellular components prior to death.

A key question for future research is how the negative response to dead cells in terms of population growth in the high-nutrient environment has evolved. One possible explanation is an adaptive response that might protect bacteria from stress. Dead cells, especially dead conspecifics, may be reliable indicators of lethal stress nearby (12,13). Recent work found evidence of such ‘necrosignalling’ in *E. coli*, by showing that swarming *E. coli* cells respond to released protein from dead clone mates, increasing their ability to withstand antibiotic stress and subsequently swarm on agar with an otherwise lethal concentration of antibiotic (15). This increase in antibiotic resistance is only observed in swarming *E. coli*, and agrees with both our transcriptomic data pointing to a link between a dead-cell response and motile behavior, and our failure to observe altered swimming behavior in the presence of dead cells. Our work also goes beyond these past findings by addressing the effect of dead cells on population growth, and how this depends on local nutrient status. We speculate the increased transcription of motility-associated genes may reflect a ‘fleeing’ response to dead cells and associated stressors. Future work testing whether the overexpression of motility-associated genes and reduced population growth we observed in the presence of dead cells confers a fitness benefit to bacteria in other types of stressful conditions will shed light on this idea. We note also that, while our results strongly implicate motility-associated genes in the observed response to dead cells, we also observed changes in gene expression for some other types of genes. Although our genetic knockouts indicated defective motility systems ameliorated the observed effect of dead cells, we do not rule out other types of responses.

In conclusion, our results show dead bacteria influence live bacterial cultures, including a novel, growth-suppressive effect in a seasonal environment where bacteria are initially supplied with other nutrients. This depends critically on how the bacteria involved died (lysed or not) and which environment they are growing in. The response seems to be rooted in upregulation of motility-and chemotaxis-associated genes. Because many environments have a diverse range of nutrient contents and lethal stressors, these results are likely relevant in many natural conditions. For example, the nutrient medium we used (LB) mainly contains amino acids as carbon source (43), thought to be a major component of the nutrients available in the gut (44). Similarly, because of the huge abundance of bacteria in the gastrointestinal tract, dead cells are likely to be present in significant quantities in such environments, some of which may be lysed as in our experiments, due for example to killing by other bacteria, antibiotics or viruses. Another key implication of our findings is in population biology, where theoretical models of microbial populations frequently assume that dead cells simply exit the model. Our results suggest instead that dead cells can have important effects by regulating population density.

## Materials & Methods

### Organisms and growth conditions

We used various bacterial strains in different experiments (Table S3), and for most experiments we used two types of growth medium: LB (Lennox lysogeny broth, Sigma) and M9 mineral medium (containing M9 salts [33.7 mM Na_2_HPO_4_, 22.0 mM KH_2_PO_4_, 8.55 mM NaCl, 9.35 mM NH_4_Cl] plus 1 mM MgSO_4_ and 0.3 mM CaCl_2_). For one experiment we supplemented M9 with 0.2 % *w*:*v* glucose, glycerol or casamino acids. For another experiment we made nutrient-depleted versions of LB by reducing the concentration of tryptone and yeast extract. All incubation was at 37°C.

### Growth assays

In our first experiments, we measured the response of *E. coli* to dead cells (prepared as described below) by filling wells in 96-well microplates (Greiner, Kremsmünster, Austria) with either 148.5 μl of dead-cell suspension or control medium (prepared as described below), after which we inoculated each well with 1.5 μl of an independent overnight culture grown in the appropriate medium. For several experiments, we used 50 μl of dead-cell suspension with 98.5 μl of fresh medium instead; this is indicated where relevant in the main text. For all experiments, we inoculated 4-12 replicate populations (depending on the experiment). Starting ODs were measured using a spectrophotometer (Tecan NanoQuant Infinite M200 Pro), after which the microplates were incubated for 24h, before measuring their final densities. We then calculated the change in OD600 over 24h for each well (ΔOD, equal to OD600_*t=24h*_ - OD600_*t=0h*_).

For some experiments, we used a second measure of bacterial growth by counting colony forming units (CFU/mL), after plating dilution series onto LB agar plates. We used Welch’s *t*-test to determine whether average yield were affected by the addition of dead cells. We also plated one culture of each treatment level on TA (tetrazolium arabinose) agar to check for viable cells from the “dead” cultures added (see below).

### Preparation of dead cells

We tested the effects of various types of dead bacteria and corresponding control treatments. In every experiment, we confirmed that dead-cell aliquots contained no viable colony-forming units by plating 50 μl onto LB-agar and incubating overnight. Additionally, we prepared dead bacteria from a marked version of the ancestral strain, K12 MG1655 Δ*Ara*, allowing us to test for regrowth from killed cultures during the assay by plating on tetrazolium agar (45). We did not observe any Δ*Ara* colonies in any of our assays. We prepared dead bacteria in the following ways:

#### 1. Heat-killed cells

We centrifuged overnight cultures of *E. coli* at 4000 rpm for 5 minutes, after which we resuspended the cultures in the appropriate growth medium. This procedure removes all medium conditioning by prior growth and allows us to investigate the effect of dead cells directly. We then heated these cultures at 100°C for 1 hour. As a control, we used 150 μl aliquots of sterile medium subjected to the exact same procedure. We then plated 50 µL of each killed culture without dilution on LB and TA agar to check for viable cells. We observed no growth on the plates after the procedure.

#### 2. Sonicated-cell lysate

We hypothesized that the effect of dead bacteria may be stronger when they are lysed. Therefore we sonicated 5 mL cultures for 5 minutes with 1 second pulses at 80 % amplitude using a SonoPlus HD2070. Prior to sonication we resuspended cultures in fresh medium. Plating showed that sonication did not kill 100% of the population, so we then filtered (0.20 μm) each lysate after sonication. All culture tubes were kept on ice between resuspension and sonication/filtration. It would in theory be possible that the negative effect of dead-cell treatments we prepared in this way resulted from nutrient depletion after resuspension but prior to sonication. We tested for this using a control experiment including tubes that were resuspended, kept on ice and then filtered (but not sonicated); this showed no evidence of significant nutrient depletion after resuspension/before sonication (Fig. S1). In some experiments we heat-treated sonicated cultures instead of filtering them; this allowed us to test for the effects of heating vs filtration (comparing sonicated-and-filtered treatments to sonicated-and-heated), which might arise from denaturation of heat-labile enzymes in the lysate, and whether any lack of effect in heat-treated cultures was due to the effects of heating, or the lack of sonication (by comparing heated vs sonicated-and-heated).

#### 3. Phage-lysed cells

We hypothesized that lysis by bacteriophages would produce similar effects compared to lysis by sonication. We diluted overnight cultures of *E. coli* 100× into 5 mL fresh LB and grew them for ~2.5 hours before resuspending in fresh LB. Then, we added ~6×10^9^ plaque-forming units (PFU) of lytic phage T7 suspended in 100 μL and further incubated until lysis was visible (~3 hrs). We filtered the lysate (0.20 μm filter) to remove any viable cells (confirmed by plating on LB agar). Note this procedure does not remove the phages. To ensure this did not influence our subsequent assays, as the live *E. coli* here we used a strain (ECOR9, from the ECOR collection (46)) that is closely related to *E. coli* K12 but fully resistant to this phage. To control for the possibility that ECOR9 would react differently to dead-cell lysate compared to *E. coli* K-12, we also tested ECOR9 with dead-cell lysate produced by sonication and compared this to K12 under the same treatment. As above, the control treatment was prepared in the same way but without any bacterial cells.

### RNA isolation and sequencing

To test for changes in gene expression upon exposure to dead cells, we extracted total RNA from *E. coli* exposed to sonicated *E. coli* or the corresponding control treatment (three replicates each) at three different timepoints (after 5, 6.5 and 24 hours of growth). We grew *E. coli* the same way as in our growth assays described above, adding 50 μL of sonicated-and-filtered dead cells or control medium. We used 50 μL here, rather than 148.5 μL as in some of our experiments, to minimise the amount of extracellular RNA in the medium that was not from our focal culture, while still producing a clear effect of dead cells on growth. Additionally, cultures were pelleted and resuspended before RNA extraction, which we expect to remove the majority of any free-floating RNA derived from the lysate rather than contained in live cells. To obtain sufficient culture volume for RNA extraction, for each replicate we pooled six separate wells inoculated from the same overnight culture and incubated in the same conditions (we did this, rather than simply incubating larger culture volumes, to keep the culture volume and growth conditions identical to the main experiments above). We then pelleted the pooled cultures, removed supernatant, and added Qiagen RNA-Protect Bacteria Reagent. We then extracted total RNA with the SV Total RNA isolation kit. Total RNA was then DNAse treated using the Ambion Turbo DNA-Free DNAse kit (Life technologies), and ribosomal RNA was removed using the MICROBExpress bacterial mRNA enrichment kit (Thermo Fisher). Resulting read-alignment was performed with bowtie2 (47) using the Ensembl genome build K12_MG1655_ASM584v2 as reference. Gene expression values were computed with the function featureCounts from the R package Rsubread (48) (using the following settings: min mapping quality 10, min feature overlap 10bp, count multi-mapping reads, count only primary alignments, count reads also if they overlap multiple genes). We computed differential expression using the generalized linear model implemented in the Bioconductor package DESeq2 (49). Genes were considered expressed if they had at least 10 reads assigned.

### Keio knockout and ASKA overexpression assays

From gene expression data above, we selected a subset of genes for further investigation. We used the omics dashboard on the Ecocyc website (50) to identify genes meeting the following criteria: (1) a fold-change of >2 at one or more timepoints, (2) more than 10 downstream genes (in the case of regulatory genes) with fold changes of >2 at at least one timepoint (downstream genes were also identified with the omics dashboard). Of the 330 genes that met at least one of these criteria, we then picked 9 genes for further investigation, prioritizing those with regulatory functions. We then took single gene knockout mutants of these genes from the Keio collection (51) and tested their response to dead cells for an altered response compared to the ancestor, using the same protocol as described earlier for sonicated, lysed cells. For two genetic knockouts that showed altered responses to dead cells compared to the wild type (*fliA* and *flhC*), we also tested for an effect of overexpression of the same genes, using strains from the ASKA library (52). Briefly, these strains each carry a plasmid containing an IPTG-inducible copy of the relevant open reading frame, allowing us to test the effect of overexpression in *E. coli*. We assayed these two strains and MG1655, as described above, both with and without 0.1 mM of IPTG to induce the plasmid-borne copies of the gene.

## Notes

### Competing Interest Statement

The authors have declared no competing interest.

## References

1. Zinser ER, Kolter R. Mutations enhancing amino acid catabolism confer a growth advantage in stationary phase. J Bacteriol. 1999 Sep;181(18):5800–7.

2. Corchero JL, Cubarsí R, Vila P, Arís A, Villaverde A. Cell lysis in Escherichia coli cultures stimulates growth and biosynthesis of recombinant proteins in surviving cells. Microbiol Res. 2001;156(1):13–8.

3. Takano S, Pawlowska BJ, Gudelj I, Yomo T, Tsuru S. Density-Dependent Recycling Promotes the Long-Term Survival of Bacterial Populations during Periods of Starvation. MBio. 2017;8(1):e02336–16.

4. Nioh I, Furusaka C. Growth of Bacteria in the Heat-Killed Cell Suspensions of the Same Bacteria. J Gen Appl Microbiol. 1968;14(4):373–85.

5. Steinhaus EA, Birkeland JM. Studies on the Life and Death of Bacteria: I. The Senescent Phase in Aging Cultures and the Probable Mechanisms Involved. J Bacteriol. 1939;38(3):249–61.

6. Zambrano MM, Kolter R. GASPing for life in stationary phase. Cell. 1996;86(2):181–4.

7. Zinser ER, Kolter R. Escherichia coli evolution during stationary phase. Res Microbiol. 2004;155(5):328–36.

8. Schink SJ, Biselli E, Ammar C, Gerland U. Death Rate of E. coli during Starvation Is Set by Maintenance Cost and Biomass Recycling. Cell Syst. 2019 Jul;9(1):64–73.e3.

9. Jakubovics NS, Shields RC, Rajarajan N, Burgess JG. Life after death: The critical role of extracellular DNA in microbial biofilms. Lett Appl Microbiol. 2013;57(6):467–75.

10. Schwechheimer C, Kuehn MJ. Outer-membrane vesicles from Gram-negative bacteria: Biogenesis and functions. Nat Rev Microbiol. 2015;13(10):605–19.

11. Turnbull L, Toyofuku M, Hynen AL, Kurosawa M, Pessi G, Petty NK, et al. Explosive cell lysis as a mechanism for the biogenesis of bacterial membrane vesicles and biofilms. Nat Commun. 2016;7.

12. Westhoff S, van Wezel G, Rozen D. Distance dependent danger responses in bacteria. Curr Opin Microbiol. 2017;in press:95–101.

13. LeRoux M, Peterson SB, Mougous JD. Bacterial danger sensing. J Mol Biol. 2015 Nov;427(23):3744–53.

14. Le Roux M, Kirkpatrick RL, Montauti EI, Tran BQ, Brook Peterson S, Harding BN, et al. Kin cell lysis is a danger signal that activates antibacterial pathways of pseudomonas aeruginosa. Elife. 2015;2015(4):1–65.

15. Bhattacharyya S, Walker DM, Harshey RM. Dead cells release a ‘necrosignal’ that activates antibiotic survival pathways in bacterial swarms. Nat Commun. 2020;11(1):1–12.

16. Finkel SE. Long-term survival during stationary phase: evolution and the GASP phenotype. NatRevMicrobiol. 2006;4(1740-1526 (Print)):113–20.

17. Farrell MJ, Finkel SE. The Growth Advantage in Stationary-Phase Phenotype Conferred by rpoS Mutations Is Dependent on the pH and Nutrient Environment. J Bacteriol. 2003;185(24):7044–52.

18. Finkel SE, Kolter R. Evolution of microbial diversity during prolonged starvation. Proc Natl Acad Sci. 1999 Mar 30;96(7):4023–7.

19. Durand PM, Rashidi A, Michod RE. How an Organism Dies Affects the Fitness of Its Neighbors. Am Nat. 2011;177(2):224–32.

20. Herker E, Jungwirth H, Lehmann KA, Maldener C, Fröhlich K-U, Wissing S, et al. Chronological aging leads to apoptosis in yeast. J Cell Biol. 2004 Feb 16;164(4):501–7.

21. Kono H, Rock KL. How dying cells alert the immune system to danger. Nat Rev Immunol. 2008;8(4):279–89.

22. Medina CB, Mehrotra P, Arandjelovic S, Perry JSA, Guo Y, Morioka S, et al. Metabolites released from apoptotic cells act as tissue messengers. Nature. 2020;1–6.

23. Weinbauer MG. Ecology of prokaryotic viruses. FEMS Microbiol Rev. 2004;28(2):127–81.

24. Lederberg J. Note: mechanism of action of penicillin. Univ Wisconsin, Dep Genet. 1956 Jan;73(1):1956.

25. Yu Z, Qin W, Lin J, Fang S, Qiu J. Antibacterial Mechanisms of Polymyxin and Bacterial Resistance. Biomed Res Int. 2015 Jan 15;2015:1–11.

26. Alteri CJ, Mobley HLT. The Versatile Type VI Secretion System. In: Virulence Mechanisms of Bacterial Pathogens, Fifth Edition. American Society of Microbiology; 2016. p. 337–56.

27. Bassler BL. How bacteria talk to each other: Regulation of gene expression by quorum sensing. Vol. 2, Current Opinion in Microbiology. Elsevier Current Trends; 1999. p. 582–7.

28. Lazazzera BA. Quorum sensing and starvation: Signals for entry into stationary phase. Vol. 3, Current Opinion in Microbiology. Elsevier Current Trends; 2000. p. 177–82.

29. Liu X, Matsumura P. Differential regulation of multiple overlapping promoters in flagellar class II operons in Escherichia coli. Mol Microbiol. 1996;21(3):613–20.

30. Liu X, Matsumura P. An alternative sigma factor controls transcription of flagellar class-III operons in Escherichia coli: gene sequence, overproduction, purification and characterization. Gene. 1995;164(1):81–4.

31. Arnosti DN, Chamberlin MJ. Secondary sigma factor controls transcription of flagellar and chemotaxis genes in Escherichia coli. Proc Natl Acad Sci. 2006;86(3):830–4.

32. Braun V, Mahren S, Ogierman M. Regulation of the Fecl-type ECF sigma factor by transmembrane signalling. Curr Opin Microbiol. 2003;6(2):173–80.

33. Raymond KN, Dertz EA, Kim SS. Enterobactin: An archetype for microbial iron transport. Proc Natl Acad Sci. 2003 Apr 1;100(7):3584–8.

34. Pratt LA, Kolter R. Genetic analysis of Escherichia coli biofilm formation: roles of flagella, motility, chemotaxis and type I pili. Mol Microbiol. 1998 Oct;30(2):285–93.

35. Wood TK, González Barrios AF, Herzberg M, Lee J. Motility influences biofilm architecture in Escherichia coli. Appl Microbiol Biotechnol. 2006;72(2):361–7.

36. Kearns DB. A field guide to bacterial swarming motility. Nat Rev Microbiol. 2010 Sep 9;8(9):634–44.

37. Garbeva P, de Boer W. Inter-specific interactions between carbon-limited soil bacteria affect behavior and gene expression. Microb Ecol. 2009 Jul;58(1):36–46.

38. Chen H, Venkat S, Wilson J, McGuire P, Chang AL, Gan Q, et al. Genome-wide quantification of the effect of gene overexpression on Escherichia coli growth. Genes (Basel). 2018;9(8):1–12.

39. Koch AL. Death of bacteria in growing culture. J Bacteriol. 1959 May;77(5):623–9.

40. Postgate JR, Hunter JR. The Survival of Starved Bacteria. J Appl Bacteriol. 1963 Oct 1;26(3):295–306.

41. Finkel SE, Kolter R. DNA as a Nutrient: Novel Role for Bacterial Competence Gene Homologs. J Bacteriol. 2001 Nov 1;183(21):6288–93.

42. Battesti A, Majdalani N, Gottesman S. The RpoS-Mediated General Stress Response in Escherichia coli. Annu Rev Microbiol. 2011 Oct 13;65(1):189–213.

43. Sezonov G, Joseleau-Petit D, D’Ari R. Escherichia coli physiology in Luria-Bertani broth. J Bacteriol. 2007;189(23):8746–9.

44. Alpert C, Scheel J, Engst W, Loh G, Blaut M. Adaptation of protein expression by Escherichia coli in the gastrointestinal tract of gnotobiotic mice. Environ Microbiol. 2009 Apr;11(4):751–61.

45. Lenski RE, Rose MR, Simpson SC, Tadler SC. Long-Term Experimental Evolution in Escherichia coli. I. Adaptation and Divergence During 2,000 Generations. Am Nat. 1991 Dec;138(6):1315–41.

46. Ochman H, Selander RK. Standard reference strains of Escherichia coli from natural populations. J Bacteriol. 1984;157(2):690–3.

47. Langmead B, Salzberg SL. Fast gapped-read alignment with Bowtie 2. Nat Methods. 2012 Apr 4;9(4):357–9.

48. Liao Y, Smyth GK, Shi W. The Subread aligner: fast, accurate and scalable read mapping by seed-and-vote. Nucleic Acids Res. 2013 May 1;41(10):e108–e108.

49. Love MI, Huber W, Anders S. Moderated estimation of fold change and dispersion for RNA-seq data with DESeq2. Genome Biol. 2014 Dec 5;15(12):550.

50. Paley S, Parker K, Spaulding A, Tomb JF, O’Maille P, Karp PD. The omics dashboard for interactive exploration of gene-expression data. Nucleic Acids Res. 2017;45(21):12113–24.

51. Baba T, Ara T, Hasegawa M, Takai Y, Okumura Y, Baba M, et al. Construction of Escherichia coli K-12 in-frame, single-gene knockout mutants: the Keio collection. Mol Syst Biol. 2006 Jan 21;2(1).

52. Kitagawa M, Ara T, Arifuzzaman M, Ioka-Nakamichi T, Inamoto E, Toyonaga H, et al. Complete set of ORF clones of Escherichia coli ASKA library (A complete set of E. coli K-12 ORF archive): unique resources for biological research. DNA Res. 2005;12(5):291–9.

