## Supplemental figures and tables for "Exposure to lysed bacteria can promote or inhibit growth of neighbouring live bacteria depending on local abiotic conditions"

**Supplementary Information**

***Supplementary methods***

**Motility assays:** To test the effect of dead cells on motility of *E. coli*, we used 0.2 % Tryptone agar (1% Tryptone, 0.5 % NaCl, 0.2 % agar) plates which allow bacterial swimming (1). Because *E. coli* MG1655 K-12 is known to be a fastidious swarmer on normal agar due to a lacking O-antigen (2), we chose to focus on swimming motility. As negative control, we used the *fliA* and *flhC* mutants, which are unable to swim (Fig. S11). We poured plates with 20 mL of Tryptone agar and dried them half-open in a laminar flow hood for 20 minutes, after which we either 1) spotted 20 μL of dead cells/control medium mixed with 5 μL of overnight culture resuspended in fresh LB in the centre of the plate, or 2) spotted 5 μL of resuspended overnight culture in the centre of the plate and 20 μL droplets of dead cells or control medium (LB) 3 cm either side of the inoculation point. We incubated the plates at 37°C wrapped in plastic to maintain humidity. Photos were taken after 24 hours. To analyze the plates, we used ImageJ software (3) to measure the maximal distance from the edge of the colony to the edge of the motility halo (maximal radius; Fig S11).

**Biofilm formation assay:** Motility genes are known to be involved in biofilm formation by facilitating cell adherence (4,5). Furthermore, dead cells release compounds that can be used in biofilm formation such as eDNA, proteins and vesicles (6–8). We therefore hypothesized increased biofilm production might explain both the lower population densities and the transcriptional changes observed when we treated *E. coli* with lysed cells in LB. We also included the *fliA* and the *flhC* knockout strains here, to test whether they had altered biofilm production behavior that could explain their different response to dead cells compared to the wild type. To test for biofilm formation, we grew *E. coli* as described in our normal growth assays (using a 1 : 50 live : dead ratio) and performed a crystal violet assay (4,9,10). We measured OD600 after 24 hours without shaking the plate as to not disturb the biofilm, after which we removed the supernatant. We washed all wells once with 150 μL PBS to remove planktonic cells, after which we added 175 μL of 0.1 % crystal violet (Merck) to stain cells. We incubated the plate for 10 minutes at room temperature, after which we removed the crystal violet and washed all wells 4 times with 150 μL PBS to remove excess stain. After removing the final wash, we air-dried the plate for 15 minutes in a laminar flow hood, after which we added 150 μL of 95% EtOH to solubilize the incorporated stain. We mixed all wells by pipetting up and down. The dye was allowed to solubilize for 10 minutes with the plate lid closed, after which we measured OD at 590 nm.

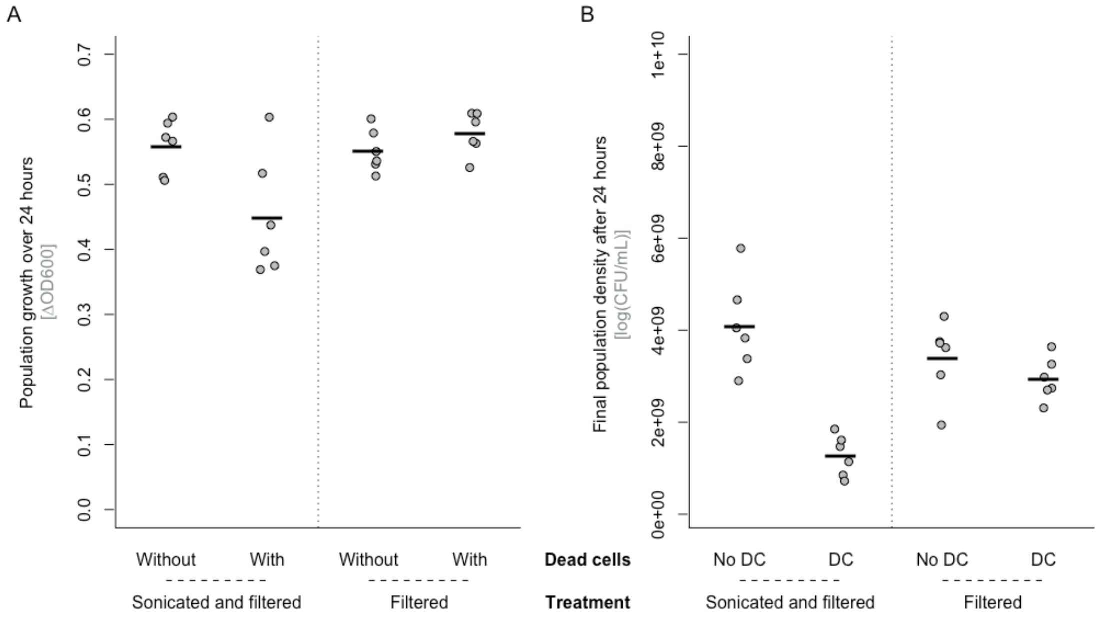

**Figure S1 Bacterial growth measured in both the change in (A) OD600 and (B) CFU/mL over 24 hours in *E. coli* in the presence and absence of either sonicated and filtered dead cells, or unsonicated filtered cells.** We prepared cells using either our standard protocol utilizing lysis by sonication and then filtration, or only filtration to control for the time cells were resuspended in fresh medium before filtration. The effect of dead cells was strongly affected by whether cells were sonicated or not (treatment × dead cells interaction, $F_{1,20} = 8.84$, $p$ = 0.008 for OD, $F_{1,20} = 22.91$, $p = 0.0001$ for CFU/mL). All points represent independent replicates (n = 5). The line shows the mean. We used a total volume of 150 µL, consisting of 1.5 µL overnight culture with 148.5 µL of dead cell suspension.

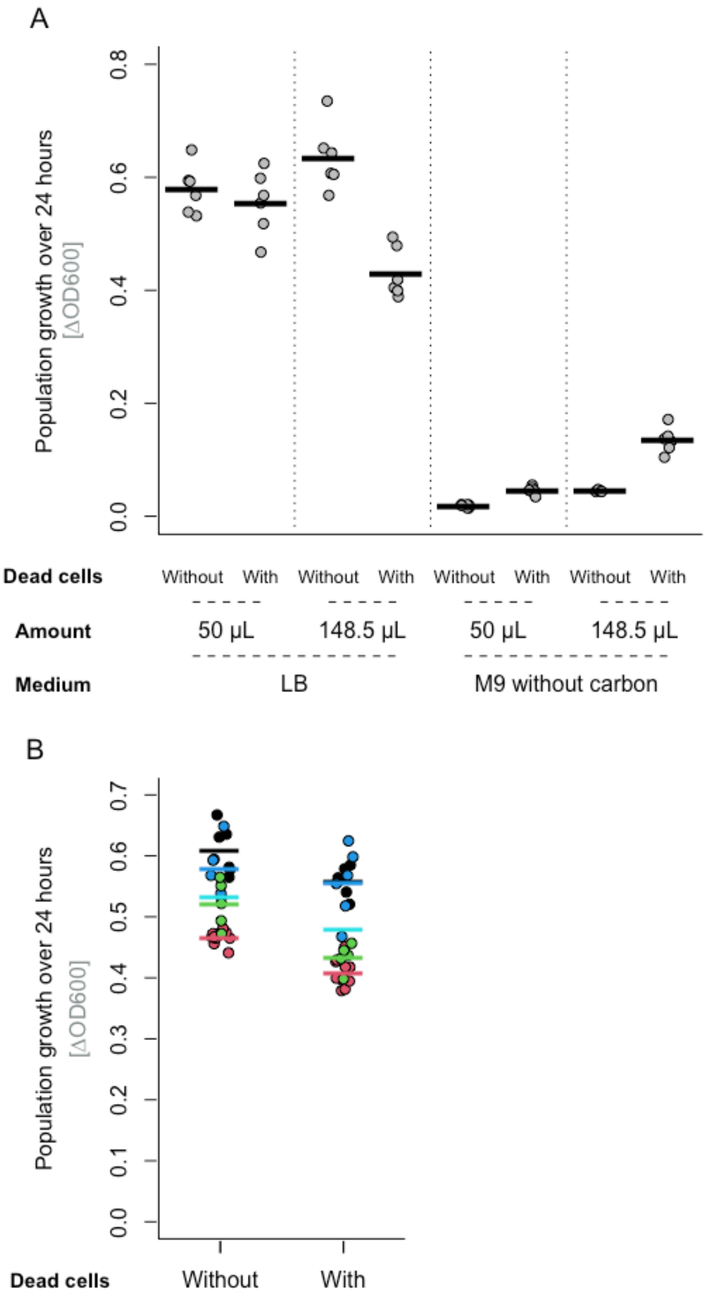

**Figure S2. Stronger effects when more dead cells are added. (A) Bacterial growth (change in optical density, OD600, over 24 hours) for *E. coli* grown in the presence/absence of different amounts of sonicated-and-filtered dead cell preparation in both nutrient-rich (LB) and minimal (M9 without carbon) medium**. We used a total culture volume of 150 µL, consisting of 1.5 µL overnight culture with either 148.5 µL of dead cell suspension, or 50 µL dead cell suspension and 98.5µL of fresh medium. All points represent independent replicates (n = 6).

Each line shows the mean for one treatment group. **(B) Effect of adding 50 µL of sonicated filtered dead cells in nutrient-rich medium (LB) across multiple experimental blocks with the same experimental design**. Different experimental blocks are coded by colours, including data from the top panel and from three other experiments using the same protocol (from figures 5, S7 and S9). Despite observing a non-significant effect in the block shown in the top panel, across the various experiments using this amount of dead cells, we observed a significant effect (effect of dead cells in two-way anova including dead cells and experimental block as factors: $F_{1,46} = 35.92$, $p < 0.0001$). This effect of dead cells was consistent across the different blocks (dead cells x dataset interaction, $F_{3,46} = 1.69$, $p = 0.18$), albeit weaker than the effect observed with 148.5 µL of dead cell preparation in the top panel.

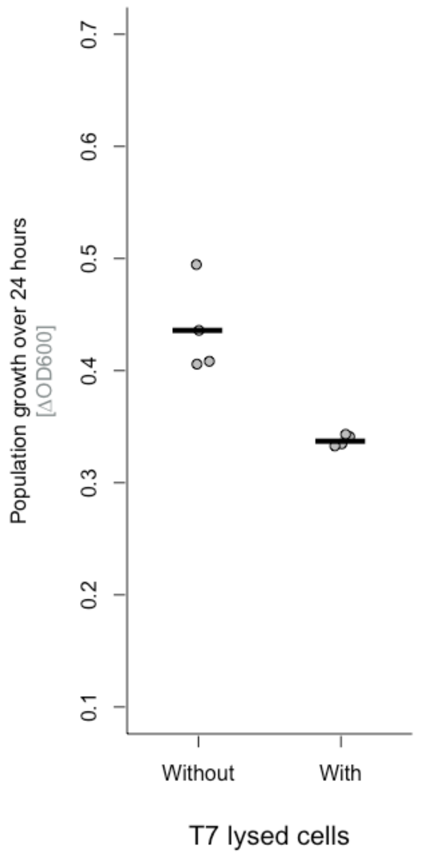

**Figure S3 Bacterial growth (change in OD600 over 24 hours) in *E. coli* in the presence and absence of T7-phage-lysed and filtered cells in nutrient-rich (LB) medium.** All points represent independent replicates (n = 4). The line shows the mean. We used a total volume of 150 µL, consisting of 1.5 µL overnight culture with 50 µL dead cell suspension and 98.5µL of fresh medium.

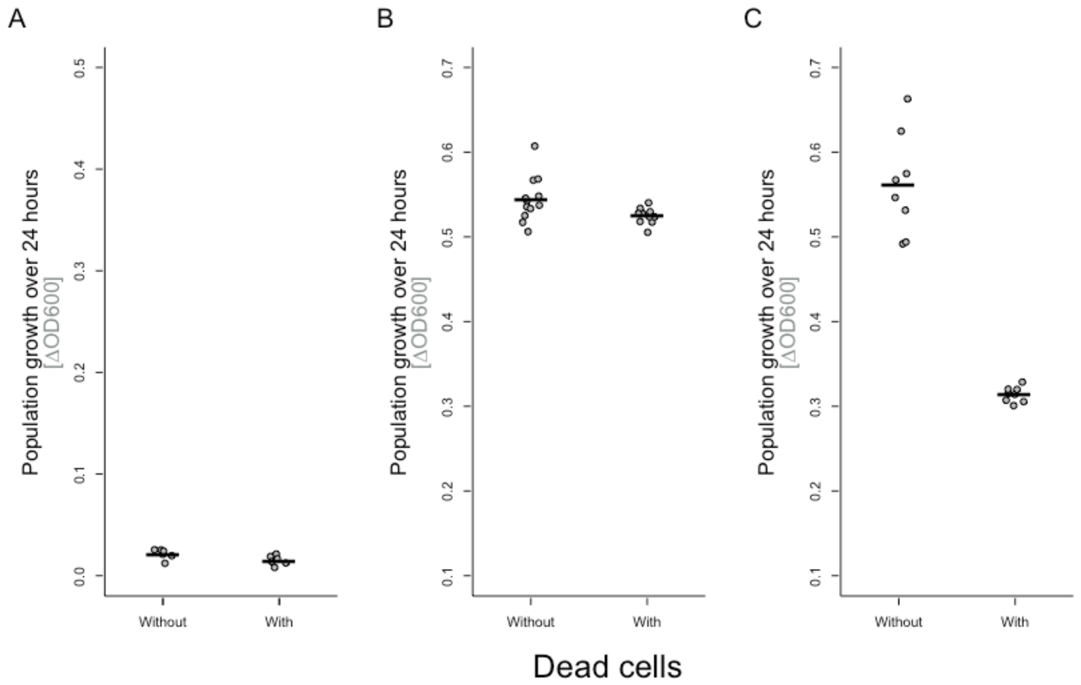

**Figure S4 Bacterial growth (change in OD600 over 24 hours) in the presence and absence of different variants of heat-killed cells in (A) minimal medium (M9, *n*=6) and (B&C) nutrient-rich medium (LB, *n*=11 and 8 respectively)**. The same type of heat-killed cells were used in (A) and (B); in (C) the dead cell preparation was first sonicated, then heat-treated. All points represent independent replicates. The line shows the mean. Note these three panels are from different experimental blocks; a direct comparison between heated and sonicated-then-heated dead cell treatments is shown in Fig. S5. We used a total culture volume of 150 µL, consisting of 1.5 µL overnight culture with 50 µL dead cell suspension and 98.5µL of fresh medium.

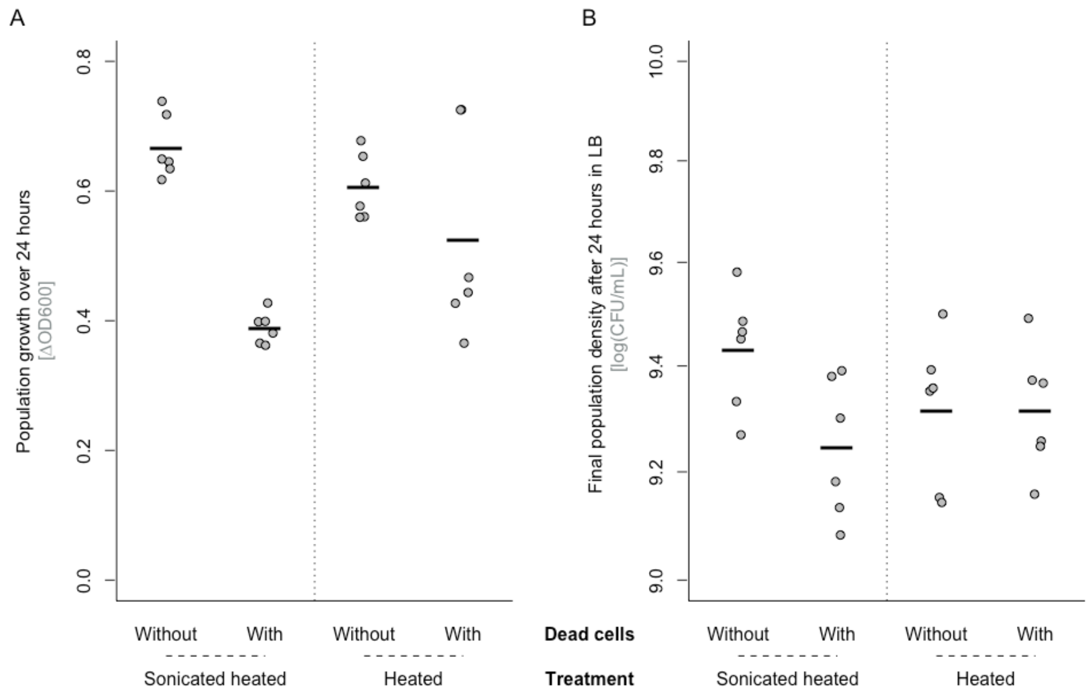

**Figure S5 Bacterial growth measured in both the change in (A) OD600 and (B) CFU/mL over 24 hours in LB in *E. coli* grown in the presence and absence of either sonicated and heated or just heated dead cells and their respective controls.** All points represent independent replicates. The line shows the mean. We used a total volume of 150 µL, consisting of 1.5 µL overnight culture with 50 µL dead cell suspension and 98.5µL of fresh medium. Note that the heated but not sonicated cells did not induce a significant effect on the population growth (Welch two-sample t-test – OD: $t = -1.20$, $df = 5.99$, $p = 0.28$; CFU/mL: $t = -0.005$, $df = 9.70$, $p = 0.99$), but the sonicated-and-heated dead cell treatment did (reported in main text).

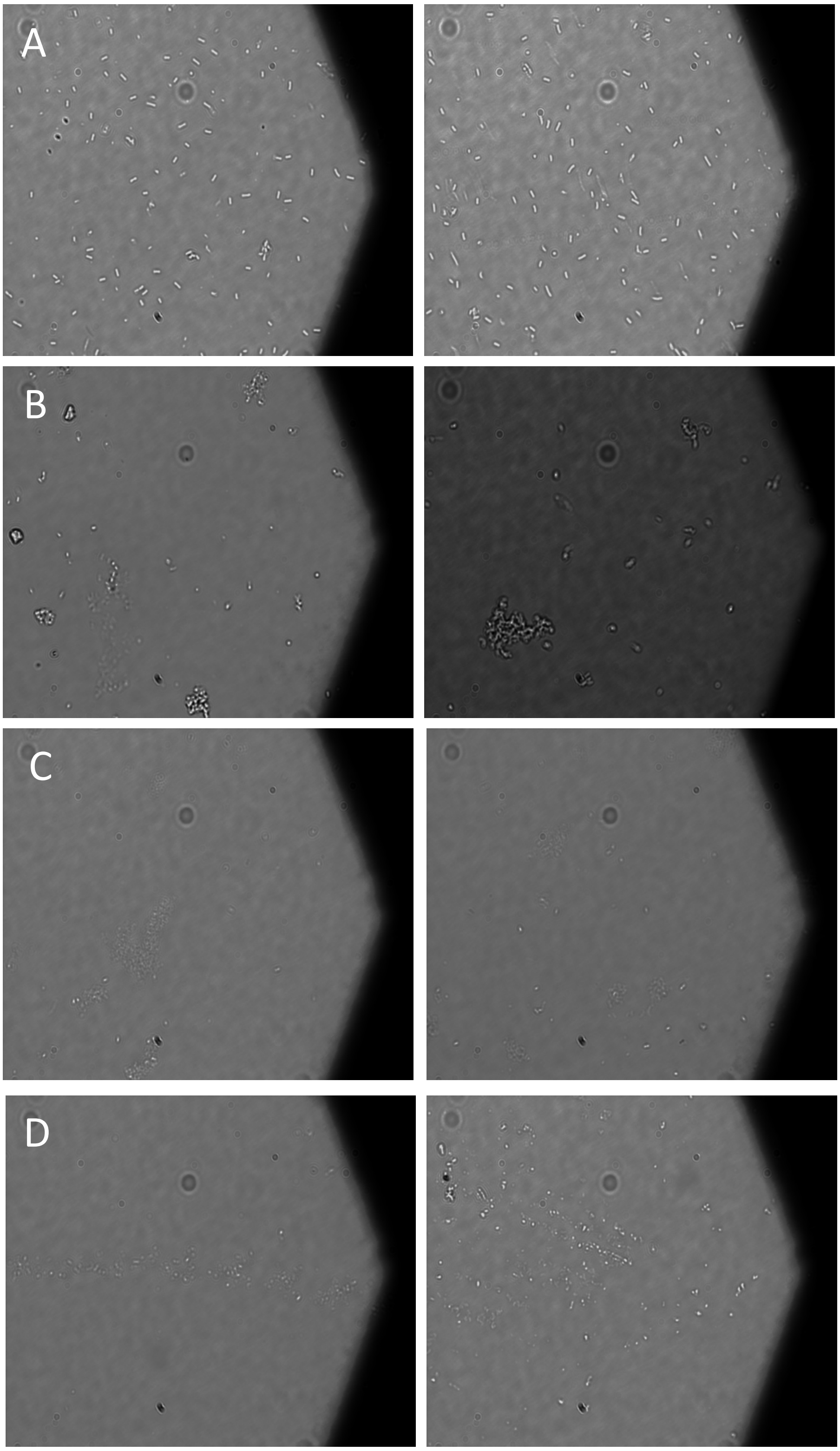

**Figure S6 Pictures of different types of live and killed cultures. (A) *E. coli* culture grown for 3 hours in LB medium (inoculated by 100× dilution from an overnight culture); (B) heat-killed cultures of *E. coli*; (C) sonicated-and-filtered cultures of *E. coli*; (D) phage-lysed and filtered cultures of *E.coli*.** Pictures were made using a Nikon Eclipse i90 microscope, using phase contrast settings on 50× magnification. The two panels in each row show independently prepared cultures.

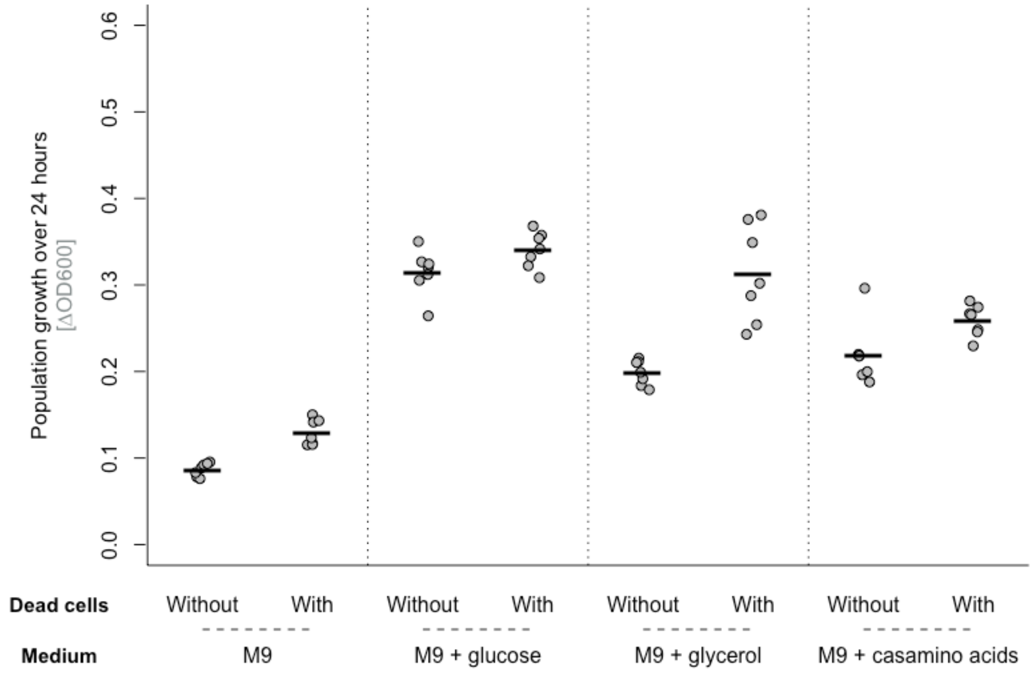

**Figure S7 Bacterial growth (change in OD600 over 24 hours) in *E. coli* in M9 medium with different carbon sources (CA= Casamino acids) in the presence and absence of sonicated filtered dead cells or control medium.** All carbon sources were added at 0.2% (*w*:*v*). All points represent independent replicates (n = 7). The line shows the mean. Note that in each of these conditions, the final bacterial density in the absence of dead cells was similar to that in relatively diluted LB treatments in Fig. 2 (OD ~0.1-0.3), where we also observed either no or weak positive responses to dead cells. Therefore, we do not exclude that populations growing to similar densities but in different abiotic conditions respond in similar ways. We used a total volume of 150 µL, consisting of 1.5 µL overnight culture with 148.5 µL of dead cell suspension.

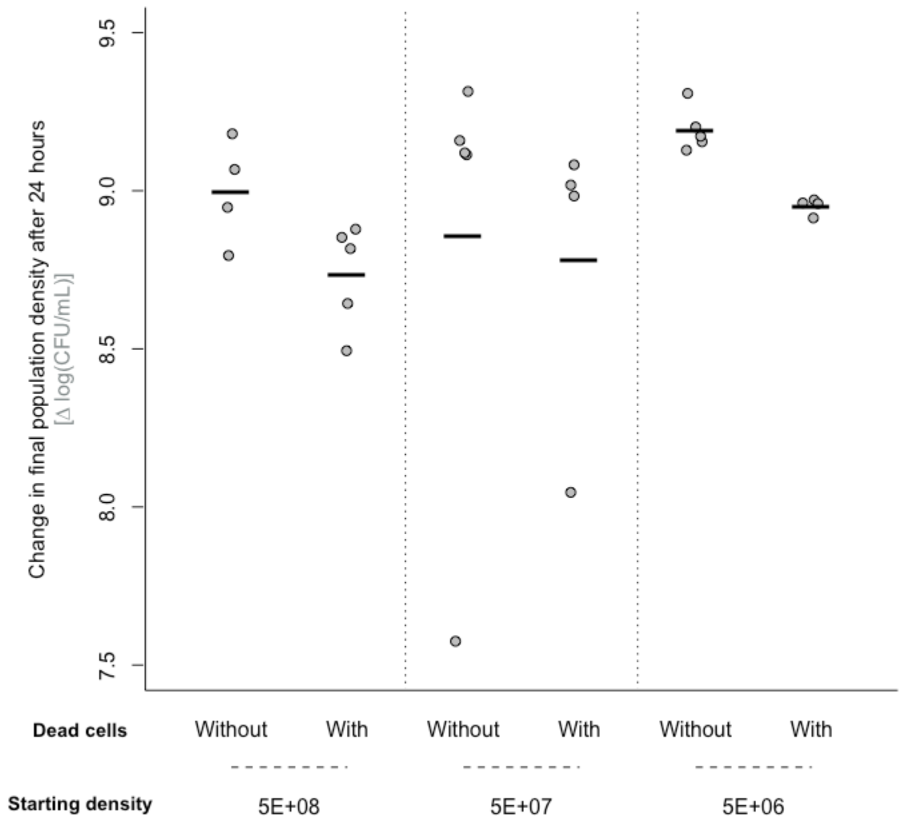

**Fig. S8 Bacterial population growth (change in CFU/mL over 24 hours in nutrient-rich medium, LB) in the presence and absence of sonicated-and-filtered dead cell preparation or control medium, using different starting densities of the live cell culture.** In all other assays, 5e+06 was used as the starting density. All points represent independent replicates (n = 5; one replicate is missing from each of the 5E+08/without, 5E+07/with and 5E+06/with treatment levels, because no colonies grew on these plates [cultures were plated at a single dilution factor]; these cultures are not plotted). Black lines show the mean. We used a total culture volume of 150 µL here, consisting of 1.5 µL overnight culture plus 148.5 µL of dead cell suspension.

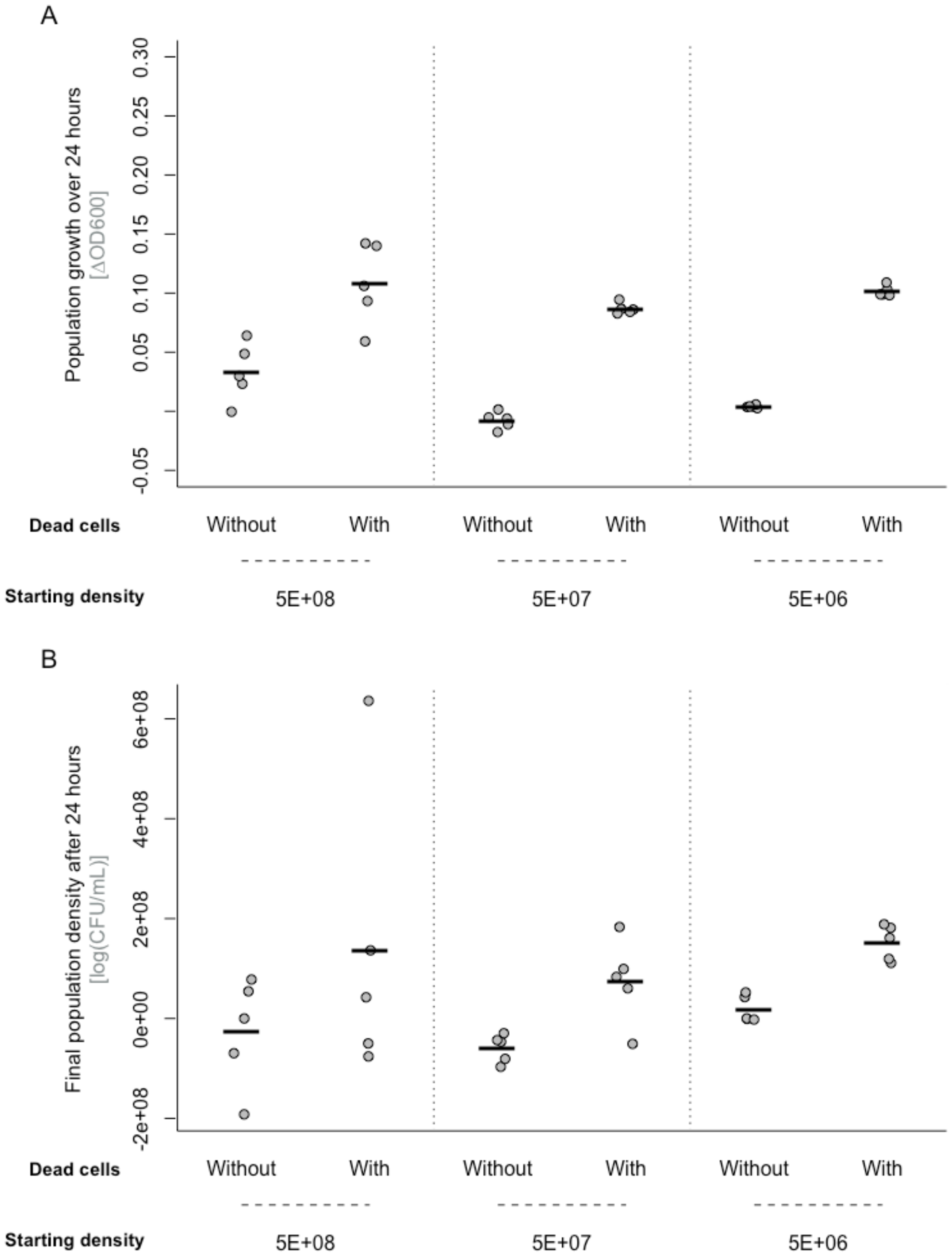

**Figure S9 Bacterial growth measured as the change in (A) OD600 or (B) CFU/mL over 24 hours in minimal medium (M9 with no additional carbon source) in the presence and absence of sonicated-and-filtered dead cell preparation using different inoculum sizes.** In all other assays, 5E+06 is used as standard inoculum. All points represent independent replicates (n =

5). The line shows the mean. We used a total volume of 150 µL, consisting of 1.5 µL overnight culture with 148.5 µL of dead cell suspension.

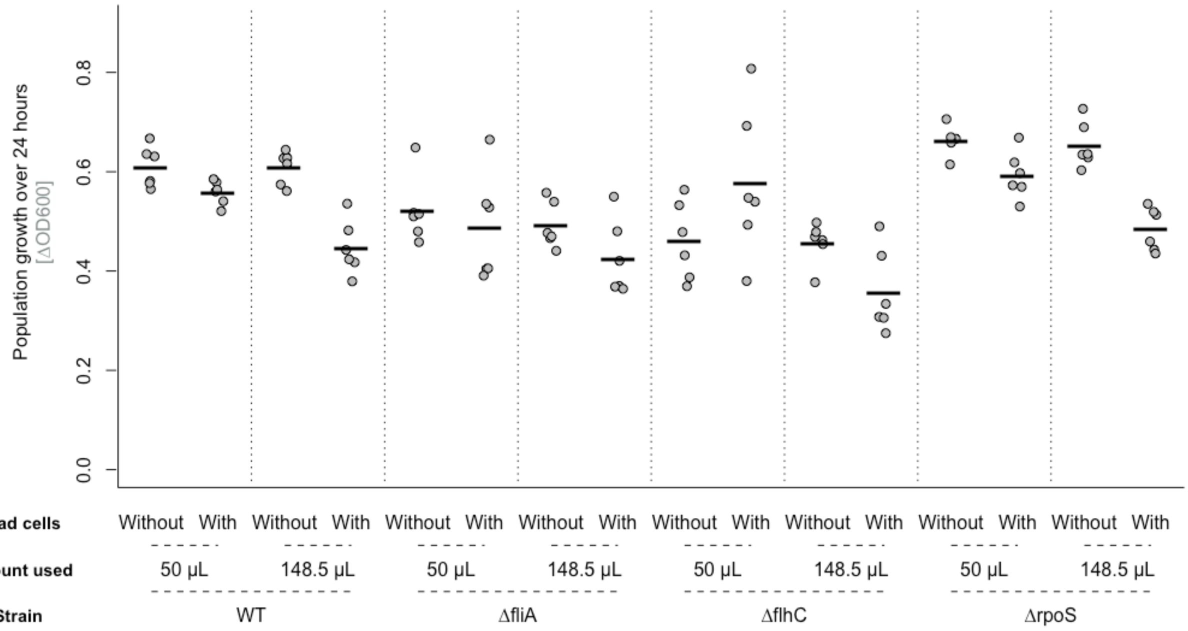

**Figure S10 Bacterial growth (change in OD600 over 24 hours) of three single-gene deletion mutants of *E. coli* BW 21533 in the presence and absence of sonicated-and-filtered dead cell preparation in LB medium, using different amounts of sonicated lysed cells**. We used a total volume of 150 µL, consisting of 1.5 µL overnight culture with either 148.5 µL of dead cell suspension, or 50 µL dead cell suspension and 98.5µL of fresh medium. All points represent independent replicates (*n* = 6). The line shows the mean.

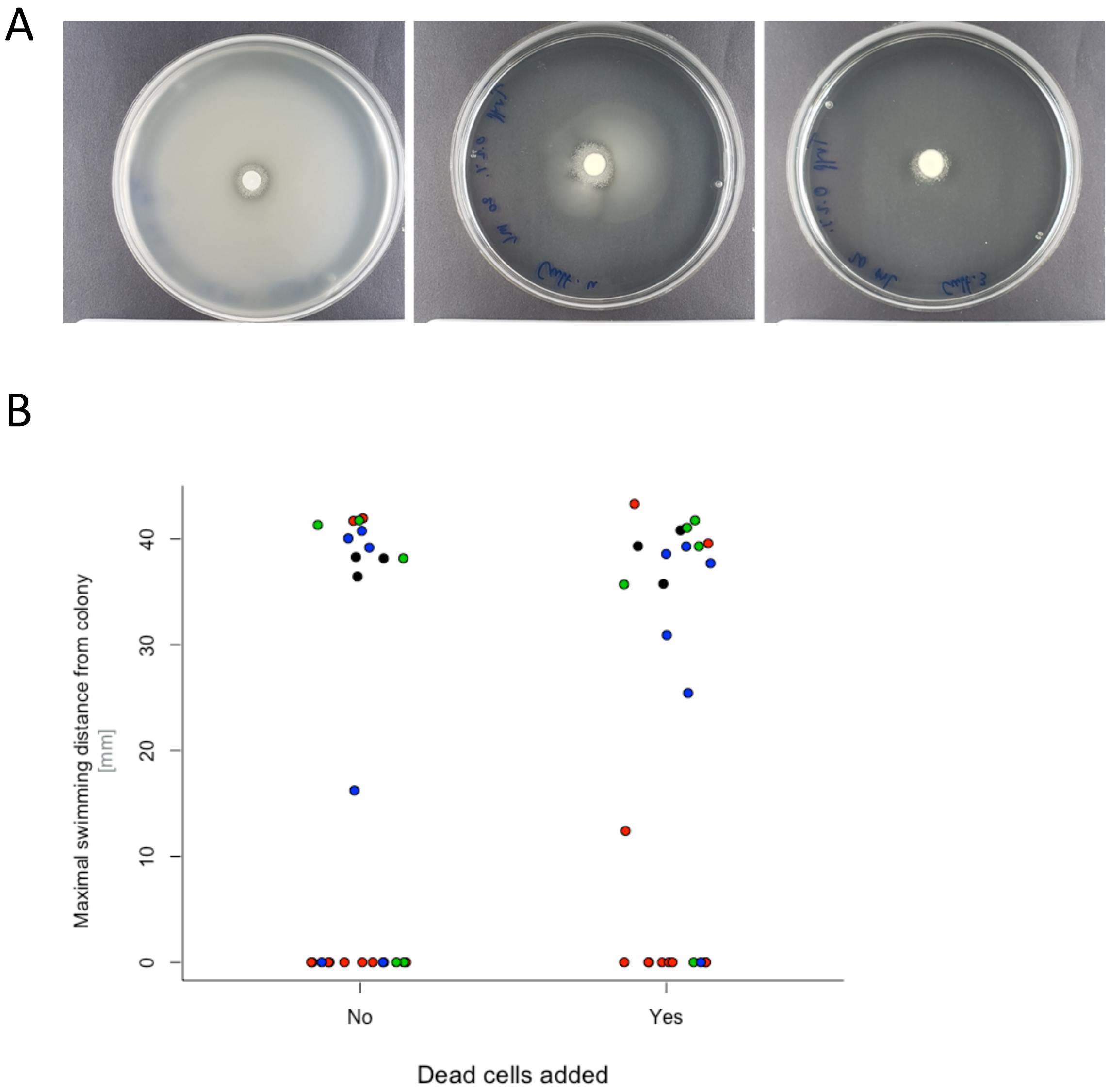

**Figure S11 No consistently visible effect of dead cells on *E. coli* swimming motility.** (A) Examples of motility plates showing different degrees of swimming, with maximal swimming on the left, partial swimming in the middle and no swimming on the right. (B) The effect of sonicated filtered dead cells on maximal swimming distance of *E. coli* on the same soft LB agar plates (0.2% agar). Each point shows one motility plate, measured in one of four different experimental blocks (different blocks shown by different colours). Maximal swimming distance on each plate is defined as the longest straight distance from the edge of the inoculated colony to the edge of the motility halo. The maximal distance possible is 45 mm (experiments were done in 90 mm petri dishes). Suspending *E. coli* in dead cells had no significant effect on average swimming distance here (linear mixed effects model with experimental block as a random factor, $t = 0.74$, $p = 0.46$). This is unchanged when we exclude plates that showed no swimming (maximum radius < 1 mm, which may result from plate-to-plate variation of agar consistency; linear mixed effects model with experimental block as a random factor, $t = -0.61$, $p = 0.55$). We also used an alternative protocol where we inoculated *E. coli* in the centre of the plate with drops of sonicated dead cells or sonicated LB (control) equally distant from the plate centre. Across 6 experimental blocks we tested 70 such plates, of which 38 showed full or partial swimming. Of these 38 plates, 2 showed an asymetric motility halo away from dead cells, but we otherwise observed no consistent bias towards or away from dead cells.

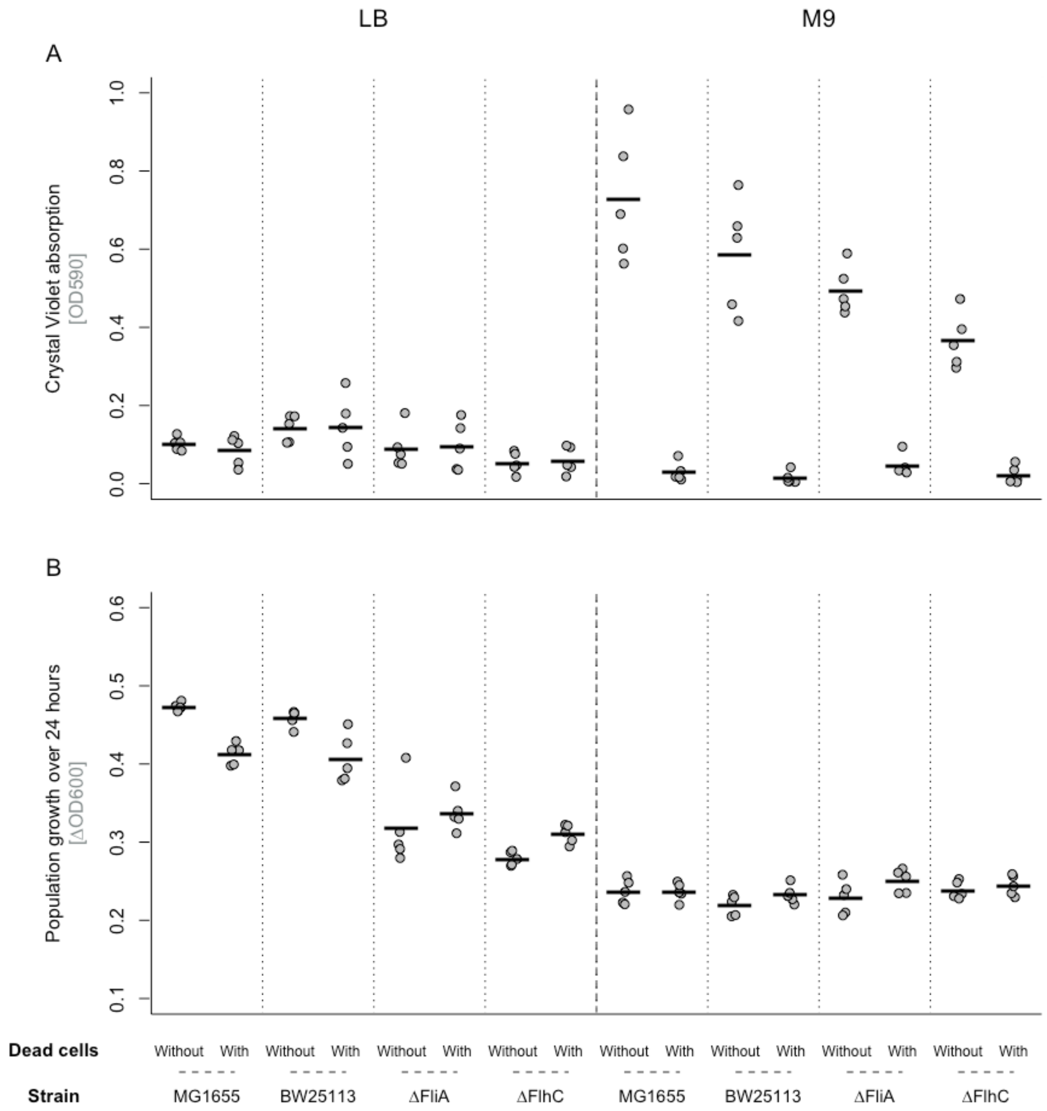

**Figure S12 Biofilm production and final density of *E. coli* MG1655, BW25113 and the *fliA* and *flhC* mutants in the presence and absence of sonicated filtered dead cells (sonicated and filtered) in LB and M9+glucose medium.** All points represent independent replicates (*n* = 5). The line shows the mean. Biofilm production was measured by crystal violet staining of adherent cells. We found no evidence of increased biofilm production in LB for either *E. coli* K12 MG1655 or BW25113 (Welch two-sample t-test - MG1655: t = 0.88, df = 5.50, p = 0.41; BW25113: t = -0.08, df = 5.42, p = 0.94), even though these cultures showed the same reduced growth yield in the presence of dead cells as we observed before (Welch two-sample t-test, t

= 9.51, df = 5.07, p = $p < 0.001$ for MG1655; t = 3.53, df = 4.94, p = 0.017 for BW25113). We used a total volume of 150 µL, consisting of 1.5 µL overnight culture *with* 50 µL dead cell suspension and 98.5µL of fresh medium.

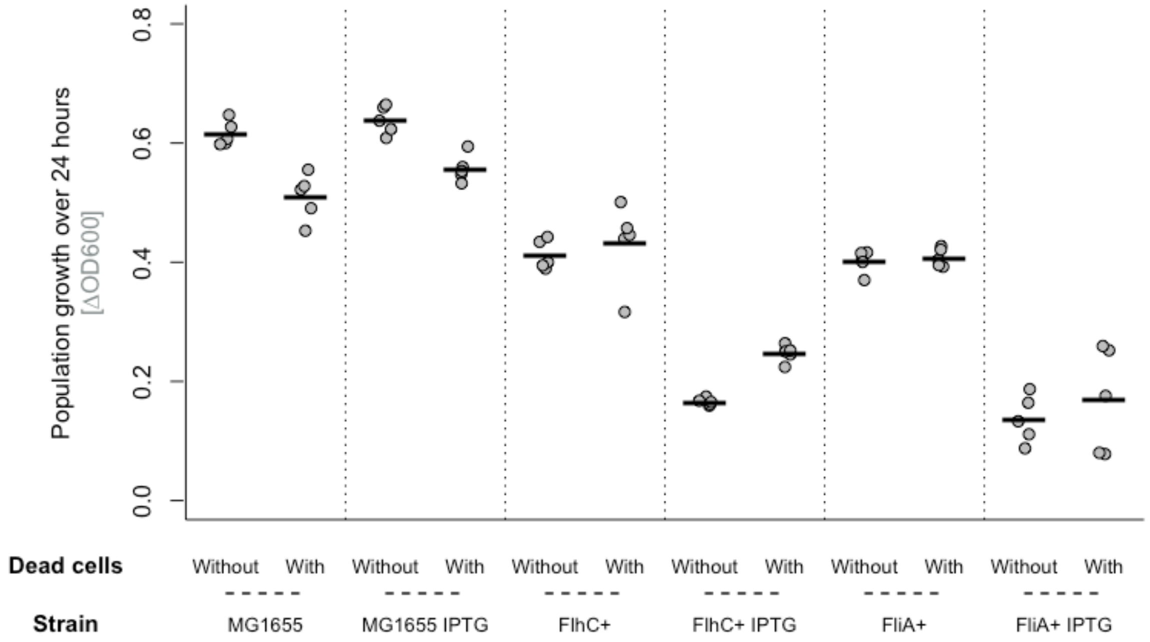

**Figure S13 Bacterial growth (change in OD600 over 24 hours) of ASKA overexpression strains of *E. coli* and the wild-type in the presence and absence of sonicated-and-filtered dead cell preparation in LB medium**. Strains were either induced or not with 0.1 mM IPTG (see *x*-axis). All points represent independent replicates ($n$ = 5). The line shows the mean. We used a total volume of 150 µL, consisting of 1.5 µL overnight culture with 50 µL dead cell suspension and 98.5µL of fresh medium.

| Gene name | Fold-change | | | FDR | | |
|---|---|---|---|---|---|---|
| | After 5 h | After 6.5 h | After 24h | After 5 h | TP 6.5 | TP 24 |
| aceA | | -2.18 | | | 7.59E-18 | |
| aceB | -2.15 | | | 3.03E-32 | | |
| aceK | | -2.31 | | | 9.81E-32 | |
| acrR | | -2.54 | | | 6.24E-04 | |
| acs | | -3.50 | | | 6.27E-24 | |
| actP | | -3.50 | | | 4.76E-19 | |
| adiY | | 2.49 | | | 1.16E-03 | |
| afuC | 2.15 | | | 3.30E-02 | | |
| aldA | | -2.16 | | | 1.46E-16 | |
| aldB | -2.05 | | | 4.14E-07 | | |
| ansP | | -2.12 | | | 8.14E-03 | |
| argB | | 2.72 | | | 3.73E-04 | |
| argC | | 2.23 | | | 1.49E-02 | |
| argH | | 2.48 | | | 6.90E-09 | |
| argI | | 4.65 | | | 1.52E-02 | |
| arpA | 2.10 | | | 8.22E-02 | | |
| artJ | | 2.10 | | | 1.25E-02 | |
| aslB | 2.16 | | | 5.85E-02 | | |
| asnA | 2.89 | | | 1.80E-27 | | |
| asnB | 2.72 | | | 1.16E-32 | | |

| | | | | | | |
|---|---|---|---|---|---|---|
| astA | | -2.71 | | | 7.86E-05 | |
| astD | | -2.34 | | | 4.80E-05 | |
| astE | | -2.54 | | | 1.32E-02 | |
| azuC | | 3.21 | | | 1.82E-02 | |
| bfd | | 3.47 | | | 8.24E-10 | |
| blc | -2.28 | | | 1.10E-04 | | |
| borD | | -2.38 | | | 1.03E-02 | |
| bsmA | | -2.12 | | | 2.31E-05 | |
| **cadA** | **4.20** | **2.54** | | **4.99E-25** | **6.49E-02** | |
| cadB | 2.08 | | | 3.91E-02 | | |
| caiF | | -2.51 | | | 2.86E-03 | |
| **carA** | **-4.09** | **-3.14** | | **2.25E-21** | **2.11E-24** | |
| **carB** | **-4.25** | **-5.91** | | **3.67E-39** | **4.74E-75** | |
| chbA | | 2.11 | | 4.51E-02 | | |
| cheA | | | 6.28 | | | 1.12E-05 |
| cheB | | | 4.99 | | | 2.49E-13 |
| cheR | | | 4.82 | | | 4.70E-04 |
| cheW | | | 5.86 | | | 3.53E-04 |
| cheY | | | 5.24 | | | 1.59E-12 |
| cheZ | | | 7.36 | | | 1.44E-16 |
| citG | 3.47 | | | 2.84E-04 | | |
| coaD | | 2.09 | | | 2.90E-05 | |
| codB | | -3.10 | | | 2.00E-09 | |

| | | | | | | |
|---|---|---|---|---|---|---|
| csiE | -4.58 | | | 8.05E-16 | | |
| cspB | 2.76 | | | 4.27E-05 | | |
| cspG | 3.35 | | | 3.20E-04 | | |
| cstA | | -3.30 | | | 5.66E-35 | |
| cvpA | | -2.40 | | | 3.59E-09 | |
| cysD | | | 2.87 | | | 3.46E-02 |
| cysH | | | 2.23 | | | 5.44E-02 |
| cysP | | | 2.03 | | | 1.18E-02 |
| cysU | | -2.10 | | | 7.91E-12 | |
| cysW | | | 2.91 | | | 9.09E-04 |
| dmlA | | | | | | |
| **dmlA** | **-3.64** | **-4.74** | | **2.37E-41** | **5.12E-35** | |
| dmlR | -2.12 | | | 4.46E-04 | | |
| ***dppA*** | ***-2.13*** | ***-2.50*** | | ***6.04E-23*** | ***1.26E-32*** | |
| dppB | | -3.92 | | | 2.36E-61 | |
| dppC | | -4.16 | | | 1.05E-40 | |
| dppD | | -4.13 | | | 1.65E-37 | |
| dppF | | -3.90 | | | 5.92E-37 | |
| **entA** | **-2.30** | **-2.11** | | **9.34E-03** | **5.91E-03** | |
| **entB** | **-2.73** | **-2.44** | | **2.41E-03** | **7.27E-04** | |
| entC | | 3.24 | | | 2.70E-03 | |
| entF | | -2.33 | | | 1.72E-05 | |
| entH | -2.10 | | | 6.92E-02 | | |

| | | | | | | |
|---|---|---|---|---|---|---|
| entS | 2.90 | | | 2.00E-04 | | |
| **exbB** | **2.16** | **2.57** | | **4.86E-11** | **5.26E-27** | |
| **exbD** | **2.16** | **2.19** | | **3.84E-11** | **1.10E-11** | |
| exuT | -2.01 | | | 3.07E-04 | | |
| fadA | | -2.90 | | | 4.71E-06 | |
| fadB | | -2.52 | | | 1.61E-09 | |
| fecA | | 2.37 | | | 2.56E-08 | |
| **fecI** | **2.02** | **4.03** | | **1.37E-04** | **2.24E-15** | |
| **fecR** | **2.24** | **4.03** | | **5.13E-05** | **1.07E-10** | |
| fepB | | 2.03 | | | 2.16E-02 | |
| **fepD** | **3.19** | **2.19** | | **2.40E-03** | **3.03E-02** | |
| **fhuF** | **2.38** | **3.54** | | **4.13E-10** | **2.96E-08** | |
| fimA | 2.38 | | | | 2.13E-38 | |
| **fimC** | **2.72** | **2.41** | | **1.06E-15** | **4.68E-07** | |
| **fimD** | **2.99** | **2.00** | | **1.61E-22** | **3.92E-06** | |
| **fimI** | **2.06** | **2.04** | | **1.95E-02** | **2.70E-08** | |
| fiu | | 2.57 | | | 3.00E-03 | |
| flgA | | | 2.36 | | | 1.92E-03 |
| ***flgB*** | | ***2.17*** | ***5.86*** | | ***2.78E-02*** | ***3.27E-17*** |
| ***flgC*** | | ***2.02*** | ***6.68*** | | ***1.98E-02*** | ***5.50E-18*** |
| ***flgD*** | ***2.09*** | | ***5.50*** | ***1.89E-04*** | | ***5.12E-09*** |
| ***flgE*** | ***2.06*** | | ***4.89*** | ***2.17E-07*** | | ***2.51E-05*** |
| ***flgF*** | ***2.25*** | | ***5.58*** | ***8.29E-05*** | | ***5.69E-28*** |

| | | | | | | |
|---|---|---|---|---|---|---|
| ***flgG*** | ***2.30*** | | ***4.69*** | ***3.01E-06*** | | ***4.35E-19*** |
| ***flgH*** | | ***2.12*** | ***3.78*** | | ***1.35E-02*** | ***1.38E-11*** |
| ***flgI*** | ***2.05*** | | ***4.00*** | ***2.08E-03*** | | ***8.59E-16*** |
| flgJ | | 4.41 | | | | 2.73E-11 |
| ***flgK*** | ***2.02*** | | ***6.19*** | ***1.86E-03*** | | ***1.55E-07*** |
| flgL | | | 4.96 | | | 1.05E-04 |
| flgM | | | 3.34 | | | 2.93E-05 |
| flgN | | | 3.53 | | | 6.03E-08 |
| flhA | | | 3.01 | | | 7.93E-06 |
| ***flhB*** | ***2.43*** | | ***5.31*** | ***9.01E-02*** | | ***1.92E-06*** |
| flhC | | 2.25 | | | 3.50E-06 | |
| ***fliA*** | ***2.27*** | | ***5.10*** | ***2.92E-03*** | | ***6.52E-18*** |
| fliC | | | 8.06 | | | 3.70E-06 |
| ***fliD*** | ***2.17*** | | ***4.41*** | ***3.57E-02*** | | ***4.17E-16*** |
| fliE | | | 8.40 | | | 9.46E-06 |
| ***fliF*** | | ***2.58*** | ***5.62*** | | ***1.31E-03*** | ***7.82E-05*** |
| ***fliG*** | ***2.25*** | | ***4.56*** | ***8.76E-04*** | | ***6.76E-15*** |
| fliH | | | 4.76 | | | 2.06E-11 |
| fliI | | | 3.97 | | | 3.15E-09 |
| ***fliJ*** | ***2.99*** | | ***4.11*** | ***9.73E-02*** | | ***1.67E-03*** |
| fliK | | | 4.82 | | | 2.53E-07 |
| fliL | | | 5.03 | | | 1.13E-02 |
| **fliM** | **2.37** | | **5.43** | **1.05E-04** | | **1.73E-16** |

| | | | | | | |
|---|---|---|---|---|---|---|
| **fliN** | **2.27** | | **4.72** | **4.44E-02** | | **5.64E-05** |
| fliO | | | 5.21 | | | 8.90E-04 |
| fliP | | | 4.20 | | | 4.75E-03 |
| fliS | | | 5.10 | | | 8.57E-08 |
| fliT | | | 3.53 | | | 7.01E-04 |
| fliZ | | | 4.44 | | | 8.26E-09 |
| flxA | | | 6.87 | | | 2.69E-07 |
| folK | | 2.07 | | | 7.57E-06 | |
| gadW | | 2.79 | | | 5.15E-13 | |
| gcvP | | | 2.01 | | | 5.05E-22 |
| gcvT | | | 2.04 | | | 4.28E-14 |
| gfcB | 2.81 | | | 6.16E-02 | | |
| gfcC | 3.41 | | | 5.32E-02 | | |
| **ghxP** | | **-2.81** | **2.23** | | **4.33E-09** | **3.33E-05** |
| glnH | -2.47 | | | 1.02E-04 | | |
| glnK | -3.34 | | | 2.71E-02 | | |
| gnsA | 2.13 | | | 7.56E-03 | | |
| **grxA** | **2.86** | **3.18** | | **4.18E-10** | **2.30E-06** | |
| hcr | | -2.02 | | | 6.48E-03 | |
| hdeD | | | -2.01 | | | 1.18E-02 |
| hemF | | 2.12 | | | 1.34E-08 | |
| hisC | | -2.15 | | | 7.81E-07 | |
| hisD | | -2.05 | | | 1.14E-06 | |

| | | | | | |
|---|---|---|---|---|---|
| iap | | 2.57 | | | 5.32E-09 |
| ibpA | | 2.46 | | | 3.29E-03 |
| ibpB | | 7.00 | | | 5.05E-09 |
| ilvB | | -2.78 | | | 8.56E-31 |
| ilvL | | -2.75 | | | 3.46E-12 |
| ilvM | | -2.17 | | | 2.98E-02 |
| ilvN | | -2.65 | | | 1.09E-06 |
| ilvX | | -6.25 | | | 1.76E-05 |
| insH1 | | -2.44 | | | 1.13E-02 |
| insJ | 3.20 | | | 1.58E-12 | |
| insK | 3.53 | | | 5.63E-12 | |
| iraP | 2.47 | | | 1.67E-04 | |
| lacY | | -3.23 | | | 3.00E-03 |
| lacZ | | -2.97 | | | 8.97E-18 |
| lamB | | -3.03 | | | 1.06E-27 |
| leuA | | -2.11 | | | 6.41E-03 |
| leuC | | -2.03 | | | 8.01E-03 |
| lgoR | | -2.19 | | | 9.02E-02 |
| livF | | -2.18 | | | 4.42E-02 |
| livG | | -2.69 | | | 3.00E-03 |
| livH | | -2.93 | | | 2.84E-03 |
| livJ | | -2.03 | | | 1.46E-03 |
| livM | | -2.30 | | | 1.56E-03 |

| | | | | | | |
|---|---|---|---|---|---|---|
| lsrA | -3.96 | | | 1.23E-04 | | |
| lysA | | -20.55 | | | 9.26E-60 | |
| malE | | -3.10 | | | 4.49E-24 | |
| malF | | -5.27 | | | 2.36E-16 | |
| malG | | -3.09 | | | 5.32E-05 | |
| malK | | -5.71 | | | 5.84E-26 | |
| malM | | -3.29 | | | 1.29E-09 | |
| malT | -2.41 | | | 2.29E-21 | | |
| mdtL | | 2.35 | | | 1.41E-08 | |
| melA | | -2.04 | | | 7.02E-07 | |
| melR | -2.21 | | | 5.74E-02 | | |
| metA | -2.73 | | | 8.25E-02 | | |
| metE | | 2.52 | | | 5.32E-09 | |
| mlc | -2.08 | | | 1.53E-04 | | |
| mlrA | -2.83 | | | 7.50E-07 | | |
| ***motA*** | | ***2.63*** | ***6.23*** | | ***6.00E-02*** | ***3.53E-04*** |
| motB | | | 6.68 | | | 3.04E-05 |
| mtfA | -2.13 | | | 7.16E-05 | | |
| mutM | 2.36 | | | 1.27E-12 | | |
| nadA | | | -2.77 | | | 4.83E-10 |
| nadB | | | -2.71 | | | 6.03E-07 |
| **narK** | | **-2.11** | **-2.84** | | **1.35E-02** | **2.92E-03** |
| narY | | | -2.68 | | | 8.59E-02 |

| | | | | | | |
|---|---|---|---|---|---|---|
| nlpA | | -2.25 | | | 8.42E-04 | |
| nrdI | | 5.17 | | | 2.69E-02 | |
| nupC | | 2.25 | | | 1.67E-15 | |
| osmE | | -2.01 | | | 2.62E-13 | |
| pepE | -2.06 | | | 2.73E-06 | | |
| pgaA | | 3.32 | | | 2.38E-16 | |
| pheA | | -5.46 | | | 3.02E-60 | |
| pheM | | 2.34 | | | 1.99E-02 | |
| phnO | | 2.01 | | | 4.04E-02 | |
| phoA | 2.51 | | | 3.69E-08 | | |
| pinQ | 2.47 | | | 8.65E-02 | | |
| pinR | 3.18 | | | 2.19E-02 | | |
| pmrR | | 2.82 | | | 2.20E-03 | |
| proV | | -2.32 | | | 8.20E-05 | |
| proW | | -2.22 | | | 4.87E-03 | |
| proX | | -2.21 | | | 1.69E-04 | |
| prpB | | | -3.20 | | | 1.29E-04 |
| prpC | | | -2.51 | | | 2.90E-02 |
| purC | | | 2.66 | | | 3.26E-09 |
| purD | | | 5.58 | | | 1.17E-36 |
| purF | | | 2.41 | | | 2.49E-13 |
| purH | | | 3.12 | | | 6.89E-21 |
| purK | | | 2.39 | | | 1.12E-05 |

| | | | | | | |
|---|---|---|---|---|---|---|
| **purL** | | **-2.62** | **4.69** | | **1.04E-25** | **7.13E-71** |
| **purM** | | **-2.37** | **2.46** | | **1.57E-08** | **3.15E-09** |
| **purT** | **-3.61** | **-2.35** | **4.11** | **1.60E-10** | **2.21E-04** | **6.68E-09** |
| putA | | -2.36 | | | 9.95E-21 | |
| putP | | -2.75 | | | 2.07E-22 | |
| qorA | -2.07 | | | 4.51E-07 | | |
| **ravA** | **-2.50** | **-2.06** | | **3.61E-10** | **4.00E-07** | |
| **rbsA** | **-13.05** | **-3.48** | | **1.69E-93** | **9.50E-35** | |
| ***rbsB*** | ***-2.27*** | ***-2.31*** | | ***1.72E-26*** | ***3.56E-25*** | |
| **rbsC** | **-4.46** | **-2.58** | | **1.47E-32** | **5.61E-22** | |
| **rbsD** | **-8.60** | **-4.31** | | **2.08E-35** | **7.30E-37** | |
| rbsK | -2.16 | | | 1.44E-16 | | |
| recE | | -2.80 | | | 4.44E-04 | |
| sbp | | -2.28 | | | 5.29E-08 | |
| slp | -2.07 | | | 8.48E-04 | | |
| soxR | | 2.40 | | | 6.17E-02 | |
| soxS | | 4.40 | | | 1.96E-08 | |
| sra | -2.24 | | | 1.56E-10 | | |
| srlA | | -5.26 | | | 3.67E-05 | |
| srlB | | -11.76 | | | 2.69E-06 | |
| srlD | | -2.75 | | | 2.38E-05 | |
| srlE | | -4.76 | | | 2.45E-07 | |
| sstT | | -2.22 | | | 3.67E-17 | |

| | | | | | | |
|---|---|---|---|---|---|---|
| tap | | | 9.38 | | | 7.68E-10 |
| tar | | | 8.46 | | | 4.56E-04 |
| tauB | 5.56 | | | 3.77E-02 | | |
| tdcA | -4.13 | | | 3.82E-03 | | |
| tdcB | | 4.30 | | | 2.12E-10 | |
| tdcC | | 5.92 | | | 1.17E-66 | |
| tdcD | | 5.93 | | | 1.41E-75 | |
| tdcE | | 4.92 | | | 2.27E-64 | |
| tdcF | | 5.30 | | | 5.03E-38 | |
| tdcG | | 4.56 | | | 5.06E-29 | |
| thrA | | -12.61 | | | 3.94E-188 | |
| thrB | | -7.34 | | | 2.68E-117 | |
| thrC | | -3.97 | | | 3.83E-72 | |
| thrL | | -2.85 | | | 2.99E-13 | |
| tolR | | 2.13 | | | 1.05E-10 | |
| tonB | | 2.17 | | | 5.05E-09 | |
| tpr | | -2.45 | | | 2.50E-02 | |
| treB | | -11.01 | | | 1.02E-162 | |
| treC | | -9.09 | | | 8.43E-167 | |
| trxC | | 2.49 | | | 8.84E-09 | |
| tsgA | 2.08 | | | 4.60E-04 | | |
| tsr | | | 6.06 | | | 7.96E-12 |
| tsx | | 2.03 | | | 6.11E-14 | |

| | | | | | | |
|---|---|---|---|---|---|---|
| ttdA | | 2.69 | | | 9.85E-03 | |
| ttdB | | 3.20 | | | 2.21E-02 | |
| ttdR | | -2.23 | | | 2.13E-02 | |
| **ttdT** | **2.80** | **4.10** | | **4.30E-07** | **3.88E-02** | |
| ugd | | 2.05 | | | 8.51E-02 | |
| uhpT | 2.75 | | | 3.48E-02 | | |
| uspB | -2.45 | | | 3.84E-06 | | |
| uxaA | | -2.03 | | | 6.38E-09 | |
| uxaC | | -2.32 | | | 1.45E-09 | |
| ves | | | 2.46 | | | 2.57E-02 |
| wrbA | -2.11 | | | 5.43E-18 | | |
| **xanP** | **-4.16** | **-5.29** | **3.05** | **3.26E-04** | **7.00E-18** | **8.85e-7** |
| yaaX | | -2.50 | | | 3.56E-02 | |
| yaeF | | 2.23 | | | 2.72E-02 | |
| yagE | | -2.48 | | | 9.72E-05 | |
| yagF | | -2.01 | | | 2.80E-04 | |
| **ybaE** | **-2.41** | **-2.08** | | **5.12E-02** | **3.43E-03** | |
| ybcH | | -2.07 | | | 5.00E-02 | |
| ybdD | | -3.17 | | | 5.99E-11 | |
| ycaD | | 2.19 | | | 1.29E-03 | |
| ycfJ | | 2.56 | | | 1.61E-02 | |
| ycgB | -2.02 | | | 4.43E-11 | | |
| ycgR | | | 7.46 | | | 4.58E-06 |

| | | | | | | |
|---|---|---|---|---|---|---|
| ydcJ | | -2.56 | | | 3.33E-07 | |
| ydfU | | 2.60 | | | 8.58E-02 | |
| **ydfZ** | **-3.34** | **-2.24** | | **1.22E-08** | **1.35E-12** | |
| ydhC | | 5.13 | | | 1.78E-45 | |
| ydhY | | -3.82 | | | 1.69E-02 | |
| ydiE | | 5.42 | | | 1.82E-03 | |
| ydiY | | 2.04 | | | 4.06E-04 | |
| ydjX | -2.80 | | | 3.78E-03 | | |
| yeeD | | | 2.68 | | | 3.70E-03 |
| yeeR | | -2.20 | | | 3.86E-04 | |
| yehB | 2.92 | | | 5.86E-02 | | |
| yeiQ | | 4.72 | | | 1.30E-42 | |
| yfgH | | 2.52 | | | 1.51E-02 | |
| **yfgI** | **2.23** | **3.04** | | **1.71E-02** | **5.96E-04** | |
| yfhL | | -2.21 | | | 8.98E-02 | |
| yfiM | | 2.06 | | | 8.57E-02 | |
| ygbM | | 2.63 | | | 7.67E-02 | |
| ygcS | 2.97 | | | 8.43E-02 | | |
| ygdI | | -2.15 | | | 6.22E-05 | |
| **ygfI** | **3.13** | **2.98** | | **2.01E-09** | **1.70E-04** | |
| yhbU | -2.80 | | | 6.29E-09 | | |
| yhbV | -2.11 | | | 3.91E-05 | | |
| yhcC | -4.24 | | | 3.29E-04 | | |

| | | | | | | |
|---|---|---|---|---|---|---|
| yhcN | | 2.09 | | | 2.14E-03 | |
| yhjE | | -2.36 | | | 3.14E-02 | |
| yhjG | | | 2.16 | | | 1.36E-06 |
| yhjH | | 3.43 | 9.32 | | 4.71E-02 | 3.25E-15 |
| yhjX | | 2.08 | | | 8.12E-02 | |
| yicG | | 5.55 | | | 1.75E-11 | |
| yidZ | | 2.00 | | | 9.72E-05 | |
| yjcB | 3.99 | | | 1.27E-12 | | |
| yjcH | | -4.62 | | | 1.03E-08 | |
| yjcZ | | | 3.78 | | | 3.88E-07 |
| yjeV | | | -2.10 | | | 7.17E-02 |
| **yjfN** | **-2.25** | **-2.49** | | **4.99E-02** | **2.02E-02** | |
| ymcE | 3.50 | | | 9.51E-13 | | |
| ymdF | -2.00 | | | 3.59E-02 | | |
| yncD | | 2.22 | | | 8.27E-06 | |
| yncG | | | -2.47 | | | 2.92E-03 |
| ynfE | | -2.04 | | | 5.32E-09 | |
| **ynfK** | **-4.12** | **-2.46** | | **4.80E-06** | **1.12E-02** | |
| yohC | | -2.58 | | | 2.25E-03 | |
| yohJ | | | -2.13 | | | 7.63E-04 |
| yohK | -2.28 | | | 1.10E-04 | | |
| yohO | | 2.01 | | | 3.55E-02 | |
| yojI | | 2.32 | | | 1.85E-07 | |

| | | | | | |
|---|---|---|---|---|---|
| yqaE | -2.01 | | | 4.63E-02 | |
| zraP | | 4.44 | | | 2.78E-20 |

**Table S1 All 321 genes in *E. coli* that showed > 2 fold differential expression at one or more timepoints when cultures were treated with dead *E. coli*.** Only genes with a false discovery rate < 0.1 are shown. Bold genes were > 2 fold differentially expressed at more than one timepoint. Bold and italic genes were > 2 fold differentially expressed and motility associated (defined here as falling under one of the following five GOterms: GO:0071973 (bacterial-type flagellum-dependent cell motility); GO:0071978 (bacterial-type flagellum-dependent swarming motility); GO:0044780 (bacterial-type flagellum assembly); GO:0006935 (chemotaxis); GO:0044781 (bacterial-type flagellum organization)).

| GOterm | ID |
|---|---|
| Bacterial-type flagellum organization | GO:0044781 |
| Bacterial-type flagellum assembly | GO:0044780 |
| Bacterial-type flagellum-dependent cell motility | GO:0071973 |
| Bacterial-type flagellum-dependent swarming motility | GO:0071978 |
| Chemotaxis | GO:0006935 |
| ‘de novo’ IMP biosynthetic process | GO:0006189 |
| tricarboxylic acid cycle | GO:0006099 |
| glycine decarboxylation via glycine cleavage system | GO:0019464 |
| hydrogen sulfide biosynthetic process | GO:0070814 |
| DNA unwinding involved in DNA replication | GO:0006268 |
| tRNA wobble uridine modification | GO:0002098 |

**Table S2 All 11 GO term categories that showed a significant upregulation at at least one of the timepoints.**

| ***Species*** | ***Strain*** |
|---|---|
| *Escherichia coli* | K12 MG1655 |
| | K12 MG1655 $\Delta$*Ara* |
| | BW 25113 |
| | BW 25113 $\Delta$*soxS* |
| | BW 25113 $\Delta$*lrp* |
| | BW 25113 $\Delta$*rpoS* |
| | BW 25113 $\Delta$*hns* |
| | BW 25113 $\Delta$*fnr* |
| | BW 25113 $\Delta$*fliA* |
| | BW 25113 $\Delta$*flhC* |
| | BW 25113 $\Delta$*cheY* |
| | K12 AG1 JW1907 (pCA42N (*fliA*)) GFP- |
| | K12 AG1 JW1880 (pCA42N (*flhC*)) GFP - |
| | EcoR9 |

**Table S3. Species and strains used**

**References**

1. Kearns DB. A field guide to bacterial swarming motility. Nat Rev Microbiol. 2010 Sep 9;8(9):634–44.
2. Berg HC. Swarming Motility: It Better Be Wet. Curr Biol. 2005 Aug;15(15):R599–600.
3. Schneider CA, Rasband WS, Eliceiri KW. NIH Image to ImageJ: 25 years of image analysis. Nat Methods. 2012 Jul 28;9(7):671–5.
4. Pratt LA, Kolter R. Genetic analysis of Escherichia coli biofilm formation: roles of flagella, motility, chemotaxis and type I pili. Mol Microbiol. 1998 Oct;30(2):285–93.
5. Wood TK, González Barrios AF, Herzberg M, Lee J. Motility influences biofilm architecture in Escherichia coli. Appl Microbiol Biotechnol. 2006;72(2):361–7.
6. Feng J, Ma L, Nie J, Konkel ME, Lu X. Environmental stress-induced bacterial lysis and extracellular DNA release contribute to Campylobacter jejuni biofilm formation. Elkins CA, editor. Appl Environ Microbiol. 2018 Dec 21;84(5):1–18.
7. Turnbull L, Toyofuku M, Hynen AL, Kurosawa M, Pessi G, Petty NK, et al. Explosive cell lysis as a mechanism for the biogenesis of bacterial membrane vesicles and biofilms. Nat Commun. 2016;7.
8. Whitchurch CB. Extracellular DNA Required for Bacterial Biofilm Formation. Science (80- ). 2002 Feb 22;295(5559):1487–1487.
9. O'Toole GA, Pratt LA, Watnick PI, Newman DK, Weaver VB, Kolter R. [6] Genetic approaches to study of biofilms. In: Methods in Enzymology. 1999. p. 91–109.
10. Fletcher M. The effects of culture concentration and age, time, and temperature on bacterial attachment to polystyrene. Can J Microbiol. 1977 Jan;23(1):1–6.
